# An Erg driven transcriptional program controls B-lymphopoiesis

**DOI:** 10.1101/861542

**Authors:** Ashley P. Ng, Hannah D. Coughlan, Soroor Hediyeh-zadeh, Kira Behrens, Timothy M. Johanson, Michael Sze Yuan Low, Charles C. Bell, Omer Gilan, Yih-Chih Chan, Andrew J. Kueh, Thomas Boudier, Ladina DiRago, Craig D. Hyland, Helen Ierino, Sandra Mifsud, Elizabeth Viney, Tracy Willson, Mark A. Dawson, Rhys S. Allan, Marco J. Herold, Kelly Rogers, David M Tarlinton, Gordon K. Smyth, Melissa J. Davis, Stephen L. Nutt, Warren S. Alexander

**Author notes:** Correspondence/Lead contact Dr Ashley P. Ng, The Walter and Eliza Hall Institute of Medical Research, 1G Royal Parade, Parkville, Victoria, 3052, AUSTRALIA.

## Abstract

B-cell development is initiated by the stepwise differentiation of hematopoietic stem cells into lineage committed progenitors, ultimately generating the mature B-cells that mediate protective immunity. This highly regulated process also generates clonal immunological diversity via recombination of immunoglobulin genes. While several transcription factors that control B-cell development and V(D)J recombination have been defined, how these processes are initiated and coordinated into a precise regulatory network remains poorly understood. Here, we show that the transcription factor ETS Related Gene (*Erg*) is essential for the earliest steps in B-cell differentiation. Erg initiates a transcriptional network involving the B-cell lineage defining genes, *Ebf1* and *Pax5*, that directly promotes the expression of key genes involved in V(D)J recombination and formation of the B-cell receptor. Complementation of the Erg-deficiency with a productively rearranged immunoglobulin gene rescued B-cell development, demonstrating that Erg is an essential and exquisitely stage specific regulator of the gene regulatory network controlling B-lymphopoiesis.

## INTRODUCTION

Transcription factors are critical for controlling the expression of genes that regulate B-cell development. The importance of specific B-cell transcription factors is highlighted by the phenotype of gene knockout models. Failure of B-cell lineage specification from multi-potential progenitors occurs with deletion of *Ikzf1* ^1^ and *Spi1* (*Pu.1)* ^2^, while deletion of *Tcf3 (E2A)* ^3^ and *Foxo1* ^4^ results in failure of B-cell development from common lymphoid progenitors (CLPs). Developmental arrest at later B-cell stages is observed with deletion of *Ebf1* and *Pax5* at the pre-proB and proB stages respectively ^5 6^. This sequential pattern of developmental arrest associated with loss of gene function, along with ectopic gene complementation studies ^2^, gene expression profiling ^7^ and analysis of transcription factor binding to target genes, support models in which transcription factors are organised into hierarchical gene regulatory networks that specify B-cell lineage fate, commitment and function ^8^.

Two transcription factors that have multiple roles during B-cell development are *Ebf1*, a member of the COE family, and *Pax5,* a member of the PAX family. While Ebf1 and Pax5 have been shown to bind to gene regulatory elements of a common set of target genes in a co-dependent manner during later stages of B-lineage commitment ^9^, both manifest distinct roles during different B-cell developmental stages. Ebf1 forms an early B-cell transcriptional network with E2A and Foxo1 in CLPs that appears important in early B-cell fate determination ^10^, while during later stages of B-cell development, Ebf1 acts as a pioneer transcription factor that regulates chromatin accessibility at a subset of genes co-bound by Pax5 ^11^ as well as at the Pax5 promoter itself ^12^. Pax5 in contrast, regulates B-cell genomic organisation ^13^ including the *Immunoglobulin heavy chain (Igh)* locus during V(D)J recombination, co-operating with factors such as CTCF ^14^, as well as transactivating ^15^ and facilitating the activity of the recombinase activating gene complex ^16^.

It is unclear, however, how these various functions of Ebf1 and Pax5 are co-ordinated during different stages of B-cell development. In particular, it would be important to ensure co-ordinated *Ebf1* and *Pax5* co-expression before the pre-BCR checkpoint, such that *Ebf1* and *Pax5* co-regulated target genes required for V(D)J recombination and pre-B-cell receptor complex formation are optimally expressed ^9^.

Here we show that the ETS related gene (Erg), a member of the ETS family of transcription factors, plays this vital role in B-lymphopoiesis. Deletion of Erg from early lymphoid progenitors resulted in B-cell developmental arrest at the early pre-proB cell stage and loss of V_H_-to-DJ_H_ recombination. Gene expression profiling, DNA binding analysis and complementation studies demonstrated Erg to be a stage-specific master transcriptional regulator that lies at the apex of an Erg-dependent Ebf1 and Pax5 gene regulatory network in pre-proB cells. This co-dependent transcriptional network directly controls expression of the *Rag1/Rag2* recombinase activating genes and the *Lig4* and *Xrcc6* DNA repair genes required for V(D)J recombination, as well as expression of components of the pre-BCR complex such as *CD19, Igll1, Vpreb1* and *Vpreb2*. Taken together, we define an essential Erg-mediated transcription factor network required for regulation of *Ebf1* and *Pax5* expression that is exquisitely stage specific during B-cell development.

## RESULTS

### *Erg* is required for B-cell development

To build on prior work defining the role of the hematopoietic transcription factor *Erg* in regulation of haematopoietic stem cells (HSCs) ^17^ and megakaryocyte-erythroid specification ^18^, we sought to identify whether *Erg* played roles in other haemopoietic lineages. *Erg* expression in adult hematopoiesis was first examined by generating mice carrying the *Erg*^tm1a(KOMP)wtsi^ knock-in first reporter allele (*Erg^KI^*) (Figure 1A). Consistent with the known role for *Erg* in hematopoiesis ^17–21^, significant *LacZ* expression driven by the endogenous *Erg* promoter was observed in hematopoietic stem cells (HSCs) and multi-potential progenitor cells, as well as in granulocyte-macrophage and megakaryocyte-erythroid progenitor populations, with declining activity accompanying erythroid maturation (Figure 1B with definitions of cells examined provided in Table S1 and representative flow cytometry plots in Figure S1). In other lineages, transcription from the *Erg* locus was evident in common lymphoid (CLP), all lymphoid (ALP) and B-cell-biased lymphoid (BLP) progenitor cells, as well as in B-lineage committed pre-proB, proB and preB cells and double-negative thymic T-lymphoid cell subsets, with a reduction in transcription with later B-cell and T-cell maturation (Figure 1B,C). We confirmed these findings with RNA sequencing (RNA-seq) analysis that showed significant *Erg* RNA in pre-proB, proB and preB cells (Figure 1D). This detailed characterisation of *Erg* expression raised the possibility that *Erg* plays a stage-specific function at early developmental stages of the lymphoid lineages.

**Figure 1.**
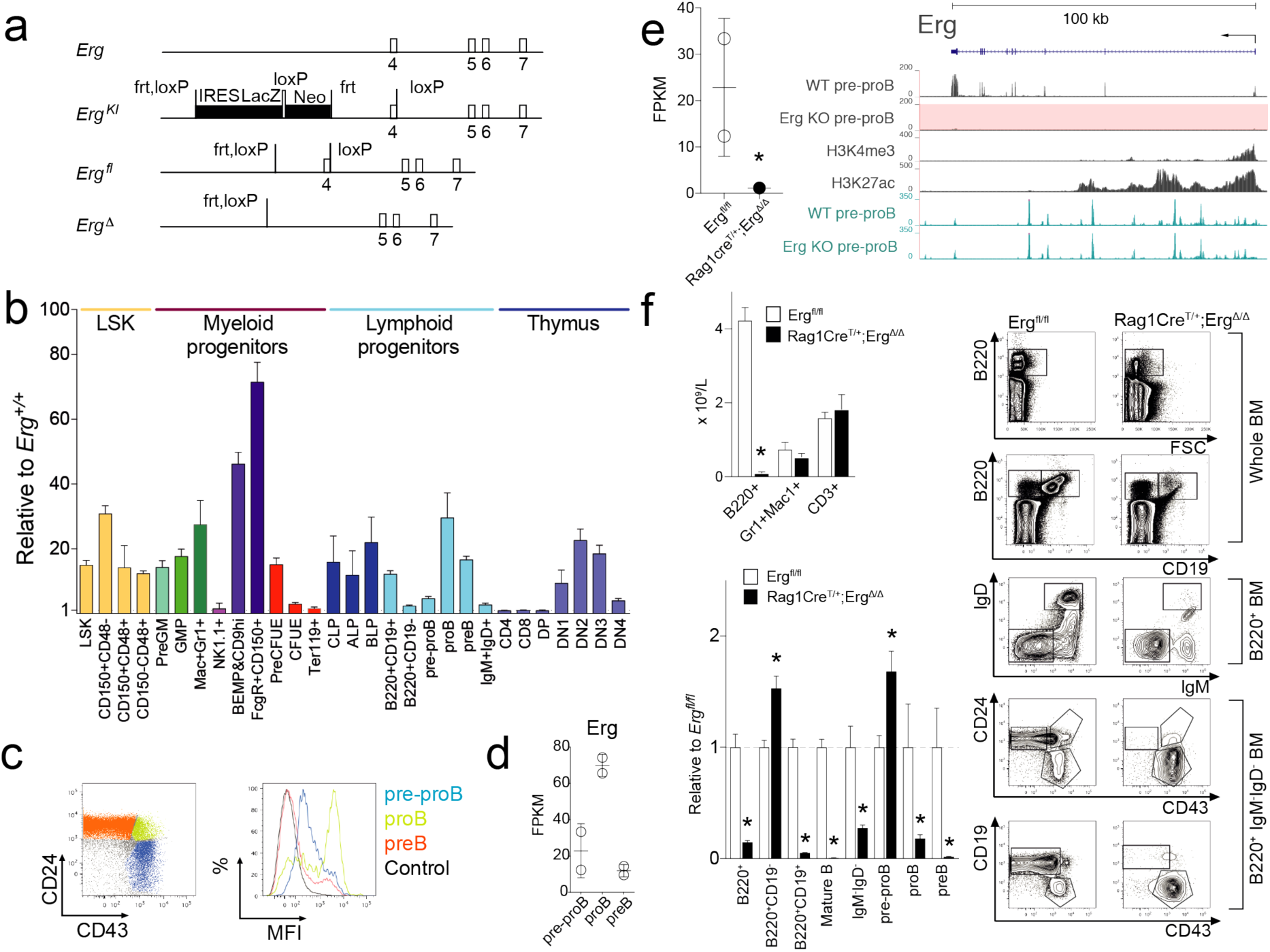
Expression and targeted disruption of *Erg* in lymphopoiesis. **A.** Wild-type (*Erg)*, *Erg*^tm1a(KOMP)wtsi^, *lacZ* reporter (*Erg^KI^*), conditional (*Erg^fl^*), and Cre recombinase-deleted (*Erg^Δ^*) alleles with exons, Cre (loxP) and Flp (frt) recombinase recognition sites. IRES, internal ribosome entry site; Neo, neomycin-resistance cassette. **B.** *Erg* transcriptional activity by *lacZ* expression in *Erg^KI^* bone marrow (BM) and thymus cell populations (see Materials and Methods and Figure S1, Table S1). Mean fluorescent intensity (MFI) ratio ± S.D of *Erg^KI^* (n=4) to C57BL/6 (n=4). *P*_adj_ < 0.027 corrected using Holm’s modification for multiple testing for each population except BM Ter119^+^ and NK1.1^+^, and thymic DP, CD4^+^CD8^-^ and CD8^+^CD4^-^ populations (*P*_adj_ > 0.05). **C.** Representative flow cytometry plots: BM pre-proB (blue), proB (green) and preB (orange) and control B220^+^IgM^-^IgD^-^ (black) (left) with *lacZ* MFI (right). **D.** *Erg* expression by RNA-seq (mean±S.D, Fragments Per Kilobase of transcript per Million mapped reads, FPKM) in *Erg^fl/fl^* pre-proB, proB and preB cells (n = 2) **E.** *Erg* RNA-seq (FPKM) in *Erg^fl/fl^* and *Rag1Cre^T/+^;Erg^Δ/Δ^* pre-proB cells (n=2) (left; *, *P* = 1.41e-5, ***Table S4***). *Erg* locus RNA-seq in *Erg^fl/fl^* (WT) and *Rag1Cre^T/+^;Erg^Δ/Δ^* (Erg KO, with pink highlighting absent expression) in pre-proB cells, H3K4me3 and H3k27ac ChIP-seq and chromatin accessibility (ATAC-seq, blue). **F.** *Erg^fl/fl^* (n=4) and *Rag1Cre^T/+;^Erg^Δ/Δ^* (n=7) B220^+^B-cell, Gr1^+^Mac1^+^myeloid-cell, and CD3^+^T-cell blood counts, mean±S.E.M; *, *P* = 6e-8 (top left). B-lymphoid populations in *Erg^fl/fl^* (n=9) and *Rag1Cre^T/+^;Erg^Δ/Δ^* (n=10) BM as ratio of cell number to *Erg^fl/fl^* (bottom left, see Table S1). *, *P*_adj_ < 0.04 corrected using Holm’s modification for multiple testing. Representative flow cytometry plots (right).

To determine whether *Erg* had a role in lymphoid development, mice carrying floxed *Erg* alleles (*Erg^fl/fl^*, Figure 1A) were interbred with *Rag1Cre* transgenic mice that efficiently delete floxed alleles in common lymphoid (CLPs) and T- and B-committed progenitor cells ^22^, but have normal lymphoid development (Figure S2A). The resulting *Rag1Cre*^T/+^*;Erg*^Δ*/*Δ^ mice specifically lack *Erg* throughout lymphopoiesis (Figure 1E, Figure S2B). While numbers of red blood cells, platelets and other white cells were normal, *Rag1Cre*^T/+^*;Erg*^Δ*/*Δ^ mice displayed a deficit in circulating lymphocytes (Table S2). This was due to a specific absence of B-cells; the numbers of circulating T-cells and thymic progenitors were not decreased (Figure 1F, Figure S2C).

B-cells are produced from bone marrow progenitor cells that progress through regulated developmental stages. B-cell development was markedly compromised in *Rag1Cre*^T/+^*;Erg*^Δ*/*Δ^ mice, with proB, preB, immature B and mature recirculating B cells (Hardy fractions C-F, defined in Table S1) markedly reduced in number or virtually absent (Figure 1F). A B-lymphoid developmental block was clearly evident at the pre-proB (Hardy fraction A-to-B) stage, with excess numbers of these cells present in the bone marrow.

### *Erg* deficient pre-proB cells have perturbed V_H_-to-DJ_H_ recombination

To further characterise the developmental B-cell block in *Rag1Cre*^T/+^*;Erg*^Δ*/*Δ^ mice, B220^+^ bone marrow progenitors were examined for *Igh* somatic recombination. Unlike cells from control *Erg^fl/fl^* mice, B220^+^ cells from *Rag1Cre*^T/+^*;Erg*^Δ*/*Δ^ mice had not undergone significant V_H_-to-DJ_H_ immunoglobulin heavy chain gene rearrangement, although D_H_-to-J_H_ recombination was relatively preserved (Figure 2A).

**Figure 2.**
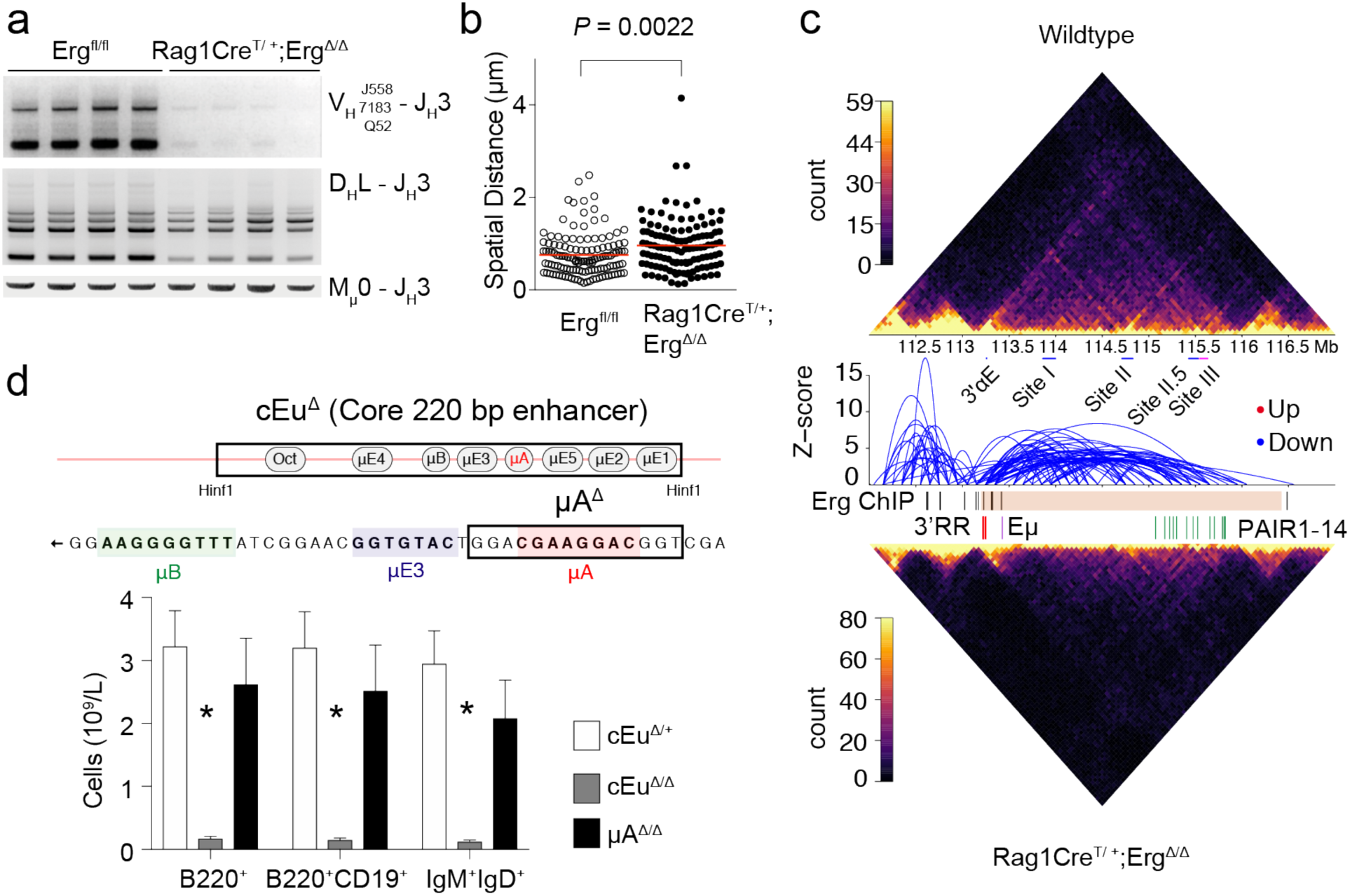
The immunoglobulin heavy chain locus in *Rag1Cre^T/+^;Erg^Δ/Δ^* mice**. A.** Genomic PCR using degenerate primers to IgH locus V_H_558, V_H_7183, V_H_Q52 segments for detection of V_H_ to DJ_H_ (top panel) and D_H_ to J_H_ (middle panel) recombination with Mu0 loading controls (bottom panel) in B220^+^ BM cells. **B.** Intra-chromosomal distance between distal VhJ558 and proximal Vh7183 V_H_ families by Fluorescent In Situ Hybridisation from (N=129) *Igh* alleles from *Erg^fl/fl^* and *Rag1Cre^T/+;^Erg^Δ/Δ^* B-cell progenitors. *P* value by Student’s two-tailed unpaired t-test. **C.** Long-range chromatin interaction by chromatin conformation and capture analysis (Hi-C) of the immunoglobulin heavy chain locus of C57BL6 (wildtype) and *Rag1Cre^T/+^;Erg^Δ/Δ^* B-cell progenitors. Reduced long-range interactions in *Rag1Cre^T/+^;Erg^Δ/Δ^* B-cell progenitors indicated by blue arcs. Erg binding by ChIP (black bars) across the heavy chain locus (pink bar) as indicated (see also ***Figure S4A***). Location of 3’regulatory region (red bars), iEμ enhancer (purple bar) and PAIR domains (green bars) are indicated. **D.** Schematic representation of iEμ enhancer with the core 220bp cEμ^Δ^ deletion and μA^Δ^ deletion shown (top). Peripheral blood counts of B220^+^, B220^+^CD19^+^ and IgM^+^IgD^+^ B-cells in cEμ^Δ/+^ (n=8), cEμ^Δ/Δ^ (n=3) and μA^Δ/Δ^ (n=7) mice (bottom). * *P* value < 0.0001 by Benjamini Hochberg correction for multiple testing. See also ***Figure S3C*.**

We next investigated the abnormalities underlying *Igh* recombination in greater detail. We first undertook fluorescence in situ hybridization (FISH) at the *Igh* locus to measure the intra-chromosomal distance between distal V_H_J558 and proximal V_H_7183 V_H_ family genes, as cell stage specific contraction of the *Igh* locus is essential for efficient V(D)J recombination ^23^. This revealed that B-cell progenitors from *Rag1Cre*^T/+^*;Erg*^Δ*/*Δ^ mice had reduced locus contraction compared to *Erg^fl/fl^* controls (Figure 2B). To assess whether other structural perturbations across the *Igh* locus were also present, chromatin conformation capture and high throughput sequencing (Hi-C) was performed. This analysis revealed a reduction of long-range interactions across the *Igh* locus in *Rag1Cre*^T/+^*;Erg*^Δ*/*Δ^ B-cell progenitors when compared to *Erg^fl/fl^* and C57BL/6 controls (Figure 2C). As these findings were also observed in *Pax5* deficient B-cell progenitors ^23 13^ reflecting a direct role for Pax5 in co-ordinating the structure of the IgH locus ^14^, we mapped *Erg* binding sites across the *Igh* locus by ChIP-seq. Unlike well-defined Pax5 binding to Pax5- and CTCF-associated intergenic regions (PAIR domains) ^14 16^, Erg binding to V_H_ families was not identified across the locus (Figure 2C, Figure S4A), suggesting that Erg was unlikely to be required structurally to maintain the multiple long-range interactions and V_H_-to-DJ_H_ recombination lacking in the *Rag1Cre*^T/+^*;Erg*^Δ*/*Δ^ B-cell progenitors. Analysis of *Igh* locus accessibility by ATAC-seq did not reveal any significant difference between *Rag1Cre*^T/+^*;Erg*^Δ*/*Δ^ pre-proB cells and control cells (***Figure S4A***) suggesting that loss of locus accessibility either by chromatin regulation ^24^ or peripheral nuclear positioning with lamina-associated domain silencing ^25^, were not mechanisms that could adequately explain reduced *Igh* locus contraction, reduction of long range interactions, and loss of V_H_-to-DJ_H_ recombination in the absence of *Erg*.

A potential role for ETS family of transcription factors in regulation of immunoglobulin gene rearrangement was proposed from experiments investigating the iEμ enhancer: a complex cis-activating element located in the intronic region between the *Igh* joining region (J_H_) and constant region (Cμ) implicated in efficient V_H_-to-DJ_H_ recombination and *Igh* chain transcription ^26^. The iEμ enhancer is proposed to nucleate a three-loop domain at the 3’ end of *Igh* interacting with the V_H_ region to juxtapose 5’ and 3’ ends of the heavy chain locus ^27^. *Erg* and its closest related ETS family member, *Fli1*, were shown to bind to the μA element and trans-activate iEμ co-operatively with a bHLH transcription factor *in vitro* ^28^. We therefore sought to determine whether the lack of Erg, and Erg binding in particular to the μA site of iEμ, could account for loss of V_H_-DJ_H_ recombination observed in *Rag1Cre*^T/+^*;Erg*^Δ*/*Δ^ mice *in vivo*. While ChIP-PCR demonstrated Erg binding to the iEμ enhancer containing the μA element (Figure S3A), mice in which the μA region (μA^Δ/Δ^) was deleted had preserved numbers of circulating mature B-cells compared to cEμ^Δ/+^ controls (Figure 2D) and intact V_H_-DJ_H_ recombination (Figure S3C). This was in contrast to cEμ^Δ/Δ^ mice, in which a core 220bp element of iEμ was deleted, that demonstrated a marked reduction of circulating mature IgM^+^IgD^+^ B-cells in peripheral blood in keeping with previous models ^29^ (Figure 2D). Together these data show that while Erg can bind to the μA region of the iEμ *in vivo*, deletion of this region did not result in significant perturbation of the B-cell development. It is therefore unlikely that Erg binding to μA element of iEμ could account for the loss of V_H_-to-DJ_H_ recombination in particular, or the *Rag1Cre*^T/+^*;Erg*^Δ*/*Δ^ phenotype in general.

### The VH10tar *IgH* knock-in allele permits B-lymphoid development in the absence of *Erg*

Given the loss of V_H_-DJ_H_ recombination associated with structural perturbation of the *Igh* locus in Erg-deficient pre-proB cells, we sought to complement the loss of formation of a functional *Igh* μ transcript and in doing so, determine whether failure to form a pre-BCR complex was a principal reason for the developmental block in *Rag1Cre*^T/+^*;Erg*^Δ*/*Δ^ mice ^30^. Complementation with a functionally re-arranged *Igh* allele in models of defective V(D)J recombination such as deletion of *Rag1, Rag2,* or components of DNA-dependent protein kinase (DNA-PK) that mediate V(D)J recombination, can overcome the pre-BCR developmental block ^31 32 33 34^.

The *IgH^VH10tar^* knock-in allele that expresses productive *Igh*^HEL^ transcripts under endogenous *Igh* locus regulation ^32^ was therefore used to generate mice that lacked *Erg* in B-cell progenitors but would undergo stage-appropriate expression of the rearranged *Igh*^HEL^ chain (*Rag1Cre*^T/+^*;Erg* ^Δ/Δ^;*IgH*^VH10tar/+^). The presence of the *IgH^VH10tar^* allele permits B-cell development in the absence of *Erg*. The bone marrow of *Rag1Cre*^T/+^*;Erg*^Δ/Δ^;*IgH*^VH10tar/+^ mice contained significant numbers of B220^+^IgM^+^ B-cells and, notably, CD25^+^CD19^+^IgM^-^ PreB cells, a population coincident with successful pre-BCR formation ^35^, that were virtually absent in *Rag1Cre*^T/+^*;Erg*^Δ/Δ^ mice (Figure 3A). Similarly, in the spleens of *Rag1Cre*^T/+^*;Erg*^Δ/Δ^;*IgH*^VH10tar/+^ mice, near normal numbers of all B-lymphoid populations were observed, in contrast to the marked reduction in *Rag1Cre*^T/+^*;Erg*^Δ/Δ^ mice (Figure 3B). Notably, IgκL chain recombination had proceeded in *Rag1Cre*^T/+^*;Erg*^Δ/Δ^;*IgH*^VH10tar/+^ cells (Figure 3C). We next tested whether the rescued *Rag1Cre*^T/+^*;Erg* ^Δ/Δ^;*IgH*^VH10tar/+^ splenic B-cells were functional in the absence of *Erg*. *Rag1Cre*^T/+^*;Erg*^Δ/Δ^;*IgH*^VH10tar/+^ splenocytes were indistinguishable from wild-type controls in *in vitro* proliferative assays using anti-μ stimulation, T-cell dependent stimulation with CD40 ligand, IL4 and IL5, or T-cell independent stimulation using lipopolysaccharide (Figure 3D). *Rag1Cre*^T/+^*;Erg* ^Δ/Δ^;*IgH*^VH10tar/+^splenic B-cells were also able to differentiate normally as measured by formation of plasma cells and IgG1 class switch recombination (Figure 3E). Circulating *Rag1Cre*^T/+^;*Erg*^Δ/Δ^*;IgH*^VH10tar*/+*^ B-cells also expressed IgD, unlike their *Rag1Cre*^T/+^;*Erg*^Δ/Δ^ counterparts (Figure 3F). These experiments demonstrated that loss of a functional *Igh* μ transcript and failure to form a pre-BCR complex was a principal reason for lack of B-cell development in *Rag1Cre*^T/+^*;Erg*^Δ*/*Δ^ mice.

**Figure 3.**
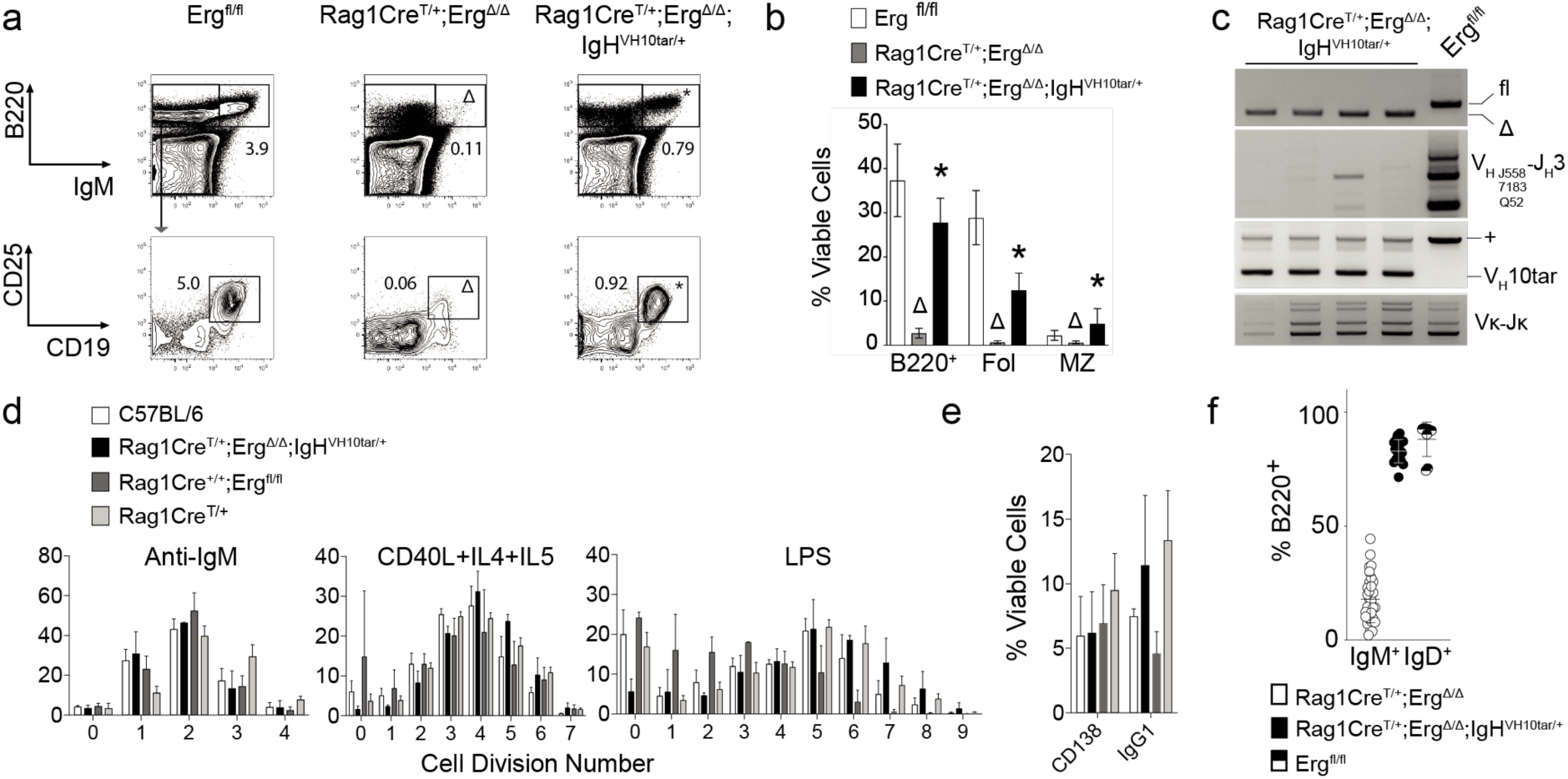
Rearranged *V_H_10_tar_* IgH allele permits *Rag1Cre*^T/+^*;Erg^Δ/Δ^* B-lymphoid development. **A.** Representative flow cytometry plots of BM B-lymphoid populations (n = 9 *Erg^fl/fl^*, n = 8 *Rag1Cre^T/+;^Erg^Δ/Δ^*, n = 8 *Rag1Cre^T/+;^Erg^Δ/Δ^;IgH^VH10tar/+^*) with mean percentage of viable cells indicated. B220/IgM profile (whole BM); CD25/CD19 profile (B220^+^IgM^-^ BM cells). Δ, *P*<10^-3^ comparing *Rag1Cre^T/+;^Erg^Δ/Δ^* to *Erg^fl/fl^*; *, *P*<0.01 comparing *Rag1Cre^T/+;^Erg^Δ/Δ^;IgH^VH10tar/+^* to *Rag1Cre^T/+;^Erg^Δ/Δ^*. **B.** Proportions of viable splenic B-lymphoid populations (n=14 *Erg^fl/fl^*, n=10 *Rag1Cre^T/+;^Erg^Δ/Δ^*, n=9 *Rag1Cre^T/+;^Erg^Δ/Δ^;IgH^VH10tar/+^*). Δ, *P*<3×10^-4^ comparing *Erg^fl/fl^* to *Rag1Cre^T/+;^Erg^Δ/Δ^*; *, *P*<2×10^-3^ comparing *Rag1Cre^T/+;^Erg^Δ/Δ^* to *Rag1Cre^T/+;^Erg^Δ / Δ^;IgH^VH10tar/+^*, mean±S.D. Fol=follicular, MZ=marginal zone (see Figure S1, Table S1). **C.** PCR of genomic DNA from B220^+^ splenocytes for *Erg* (top panel; fl, floxed allele, Δ, cre-deleted allele), V_H_-to-DJ_H_ recombination of V_H_558, V_H_7183, V_H_Q52 families (second panel), *V_H_10tar* allele (third panel) and V_κ_ light chain recombination (bottom panel). **D.** Proliferation by cell trace violet assay of wild-type (C57BL/6), *Rag1Cre^T/+;^Erg^Δ/Δ^;IgH^VH10tar/+^, Rag1Cre^+/+;^Erg^fl/fl^* and *Rag1Cre^T/+^* B220^+^ splenocytes to anti-IgM, CD40L+IL4+IL5 (T-cell-dependent) and LPS (T-cell-independent) stimulation. Mean percentage of viable cells for each cell division shown. No significant differences between genotypes were observed (*P*>0.90, 2-way ANOVA). **E.** Percentage of B220^+^ splenocytes differentiating to CD138^+^ plasma cells and undergoing IgG1 class switch recombination in response to CD40L+IL4+IL5 stimulation by flow cytometry. No significant differences were observed (*P*>0.40 for CD138 plasma cell differentiation, *P*>0.07 for IgG1 switch, corrected by Sidak’s multiple comparison test). For **D** and **E**, n=2-4 C57BL/6, 3 *Rag1Cre^T/+^;Erg^Δ/Δ^;IgH^VH10tar/+^,* 2-4 *Rag1Cre^+/+^;Erg^fl/fl^* and 3 *Rag1Cre^T/+^* mice. **F.** Percentage circulating IgM^+^IgD^+^ B220^+^B-cells in *Rag1Cre^T/+^;Erg^Δ/Δ^* (n=31), *Rag1Cre^T/+;^Erg^Δ/Δ^;IgH^VH10tar/+^* (n=17), and *Erg^fl/fl^* (n=9), *P*<10^-27^ comparing *Rag1Cre^T/+^;Erg^Δ/Δ^* to *Rag1Cre^T/+;^Erg^Δ/Δ^;IgH^VH10tar/+^* by unpaired t-test.

### Erg-deficient pre-proB cells do not express Ebf1 and Pax5 transcription factors

To define the mechanism by which *Erg* regulates V_H_-to-DJ_H_ recombination and pre-BCR formation, we undertook gene expression profiling of *Rag1Cre*^T/+^*;Erg*^Δ/Δ^ pre-proB cells. Differential gene expression and gene-ontogeny analysis of differentially expressed genes in *Rag1Cre*^T/+^*;Erg*^Δ/Δ^ pre-proB compared to *Erg^fl/fl^* pre-proB cells demonstrated deregulated expression of multiple B-cell genes (Figure 4A). These included genes encoding cell surface or adhesion receptors and core components of the pre-BCR complex *CD19, CD22, Igll1, Vpreb1, Vpreb2, CD79a and CD79b*, genes required for IgH recombination such as *Rag1* and *Rag2* and components of non-homologous end-joining repair complex associated with V(D)J recombination: *Xrcc6* (Ku70) and *Lig4*, and importantly, transcription factors implicated in B-cell development (*Ebf1, Pax5, Tcf3, Bach2, Irf4, Myc, Pou2af1, Lef1, Myb*) (Figure 4B).

**Figure 4.**
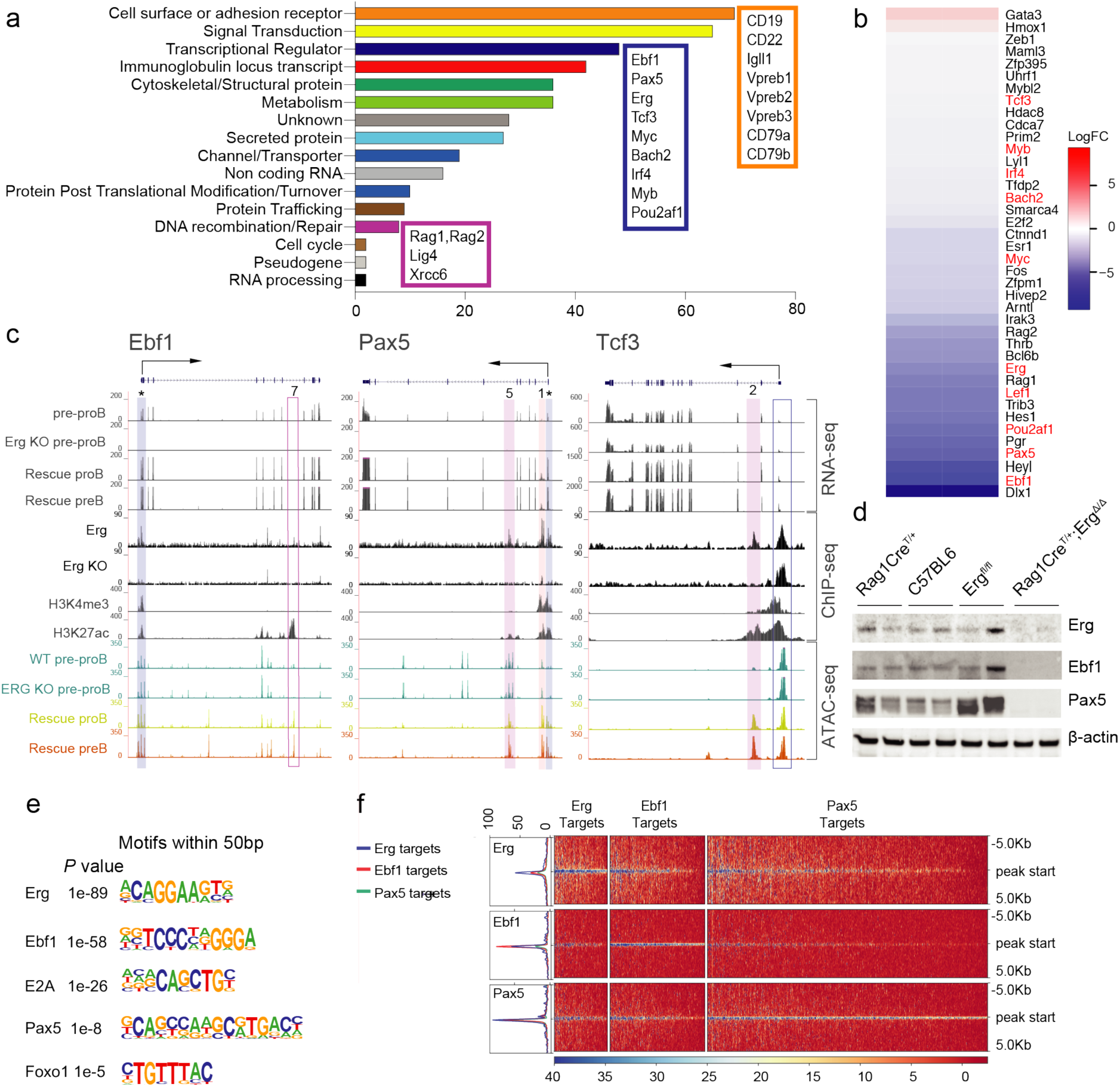
Gene expression in *Rag1Cre^T/+^;Erg^Δ/Δ^* pre-proB cells and Erg DNA binding. **A.** Differentially expressed genes in *Rag1Cre^T/+^;Erg^Δ/Δ^* pre-proB cells compared to *Erg^fl/fl^* controls, manually curated according to function based on GO term analysis (see ***Table S4***) with the number of genes for each functional category shown by the horizontal axis and selected genes highlighted in boxes. **B.** Heatmap of gene expression differences between *Erg^fl/fl^* and *Rag1Cre^T/+^;Erg^Δ/Δ^* pre-proB cells curated to transcription factors ^44^ and ordered by logFC. Selected B-cell transcription factors highlighted in red. **C.** RNA-seq for *Ebf1*, *Pax5* and *Tcf3* loci, with ChIP-seq for Erg binding in C57BL/6 B-cell progenitors and thymic *Rag1Cre^T/+;^Erg^Δ/Δ^* Erg knockout cells (Erg KO) to control for sites of non-Erg ChIP binding to DNA, H3K4me3 promoter mark, H3K27ac promoter and enhancer mark, and ATAC-seq, in *Erg^fl/fl^* pre-proB cells (pre-proB), *Rag1Cre^T/+;^Erg^Δ/Δ^* pre-proB (Erg KO pre-proB), and Erg deficient proB and preB cells in *Rag1Cre^T/+;^Erg^Δ/Δ^;IgH^VH10tar/+^*mice that develop with a functionally rearranged immunoglobulin heavy chain allele. * indicates Erg binding to the promoter region (blue bar) of *Ebf1* and *Pax5*. Solid pink bars: Erg binding to intragenic enhancer regions, with intron number as indicated. Open blue bar: promoter region with no Erg binding, open pink bar: putative enhancer region with no Erg binding. **D.** Western blot for Erg, Ebf1, Pax5 and β-actin in B-cell progenitors of genotypes indicated. **E.** Whole genome HOMER motif discovery underlying Erg bound regions in B-cell progenitors **F.** Heatmap of Erg, Ebf1 and Pax5 binding to differentially expressed genes in *Rag1Cre^T/+^;Erg^Δ/Δ^* pre-proB cells centred around the transcriptional start site (TSS) ± 5.0kB (see ***Table S5*** for all annotated ChIP binding sites).

*Ebf1* and *Pax5* are critical for B-lineage specification ^5^ and maintenance ^36, 37^ and act co-operatively to regulate a gene network in early B-cell fates ^9^. Because we observed with loss of *Erg* reduced expression of several critical B-cell genes previously identified to be controlled by *Ebf1* and/or *Pax5*, for example *CD19*, *Vpreb1*, and *Igll1* (Figure 4A), we speculated that Erg may play an important role in regulating the expression of these two essential transcription factors and their targets. To determine if *Erg* bound *Ebf1* and/or *Pax5* gene regulatory regions and directly regulated their expression, we undertook ChIP-seq analysis in wild-type B-cell progenitors and ATAC-seq to assess locus accessibility at the *Ebf1* and *Pax5* loci in the absence of Erg in *Rag1Cre*^T/+^*;Erg*^Δ/Δ^ B-cell progenitors. This demonstrated direct Erg binding to the proximal (β) promoter region of *Ebf1* ^38^ as well as to the *Pax5* promoter and *Pax5* lymphoid specific intron 5 enhancer ^12^ (Figure 4C), which together with the absence of *Ebf1*and *Pax5* expression in *Rag1Cre*^T/+^*;Erg*^Δ/Δ^ pre-proB cells and the loss of Ebf1 and Pax5 protein in *Rag1Cre*^T/+^*;Erg*^Δ/Δ^ B-cell progenitors by Western Blot, demonstrated that Erg was a direct transcriptional regulator of *Ebf1* and *Pax5* (Figure 4D). Importantly, the loss of *Ebf1* and *Pax5* expression occurred while expression of other known regulators of *Ebf1* expression, namely, *Foxo1*, *Spi1, Tcf3* and *Ikzf1* were maintained (Figure 4C ***and*** Figure S4B), and both *Ebf1* and *Pax5* loci remained accessible by ATAC-seq in *Rag1Cre*^T/+^*;Erg*^Δ/Δ^ B-cell progenitors (Figure 4C).

### A co-dependent gene regulatory network dependent on Erg, Ebf1 and Pax5

Because expression of multiple B-cell genes were deregulated in *Rag1Cre*^T/+^*;Erg*^Δ/Δ^ pre-proB cells, including those to which Ebf1 and Pax5 had been shown to directly bind and regulate, we investigated the possibility that Erg co-bound common target genes to reinforce the Ebf1 and Pax5 gene network using a genome wide motif analysis of Erg DNA binding sites in B-cell progenitors. As expected, the most highly enriched motif underlying Erg binding was the ETS motif. However, significant enrichment of *Ebf1, E2A, Pax5* and *Foxo1* binding motifs were also identified within 50bp of Erg binding sites (Figure 4E), suggesting that Erg may indeed act co-operatively with other transcription factors to regulate target gene expression in a co-dependent gene network. Analysis of the binding of each of Erg, Ebf1 and Pax5 to regulatory regions of genes that were differentially expressed in *Rag1Cre*^T/+^*;Erg*^Δ/Δ^ pre-proB cells was then undertaken. This analysis identified significant overlap of Erg, Ebf1 and Pax5 binding sites within 5kb of the transcriptional start site (TSS) of genes differentially expressed in *Rag1Cre*^T/+^*;Erg*^Δ/Δ^ pre-proB cells compared with control pre-proB cells (Figure 4F). Taken together, these data provided compelling evidence for a gene regulatory network, in which Erg is required for maintaining expression of Ebf1 and Pax5 at the pre-proB cell stage of development, as well as reinforcing expression of target genes within the network by co-operative binding and co-regulation of target genes with Ebf1 and Pax5.

To further explore our finding that Erg, Ebf1 and Pax5 form the core of a gene regulatory network in pre-proB cells, examination of Ebf1 and Pax5 binding to the *Erg* locus was undertaken. Ebf1 and Pax5 binding within intron 1 of the *Er*g locus associated with the H3K27ac mark was found, as was Pax5 binding at the *Erg* promoter (Figure S4B). To determine if Ebf1 and Pax5 directly regulate *Erg* expression, gene expression changes in B-cell progenitors from a publicly available dataset in which *Ebf1* (*Ebf1*^Δ/Δ^) or *Pax5* (*Pax5*^Δ/Δ^) had been deleted were examined (Figure 5A). Deletion of either *Ebf1* or *Pax5* resulted in reduced *Erg* expression (Figure 5B), with Ebf1 appearing to be the stronger influence. We next compared gene expression changes in *Ebf1*^Δ/Δ^ and *Pax5*^Δ/Δ^ B-cell progenitors to those genes regulated by Erg in pre-proB cells. As would be predicted if Erg, Ebf1 and Pax5 were components of a co-dependent gene regulatory network, this analysis showed a highly significant correlation in gene expression changes observed with *Ebf1* or *Pax5* deletion in B-cell progenitors and those observed with *Erg* deletion in pre-proB cells. This was noted for down-regulated genes in Erg, Ebf1 and Pax5 deficient B-cell progenitors in particular (Figure 5C).

**Figure 5.**
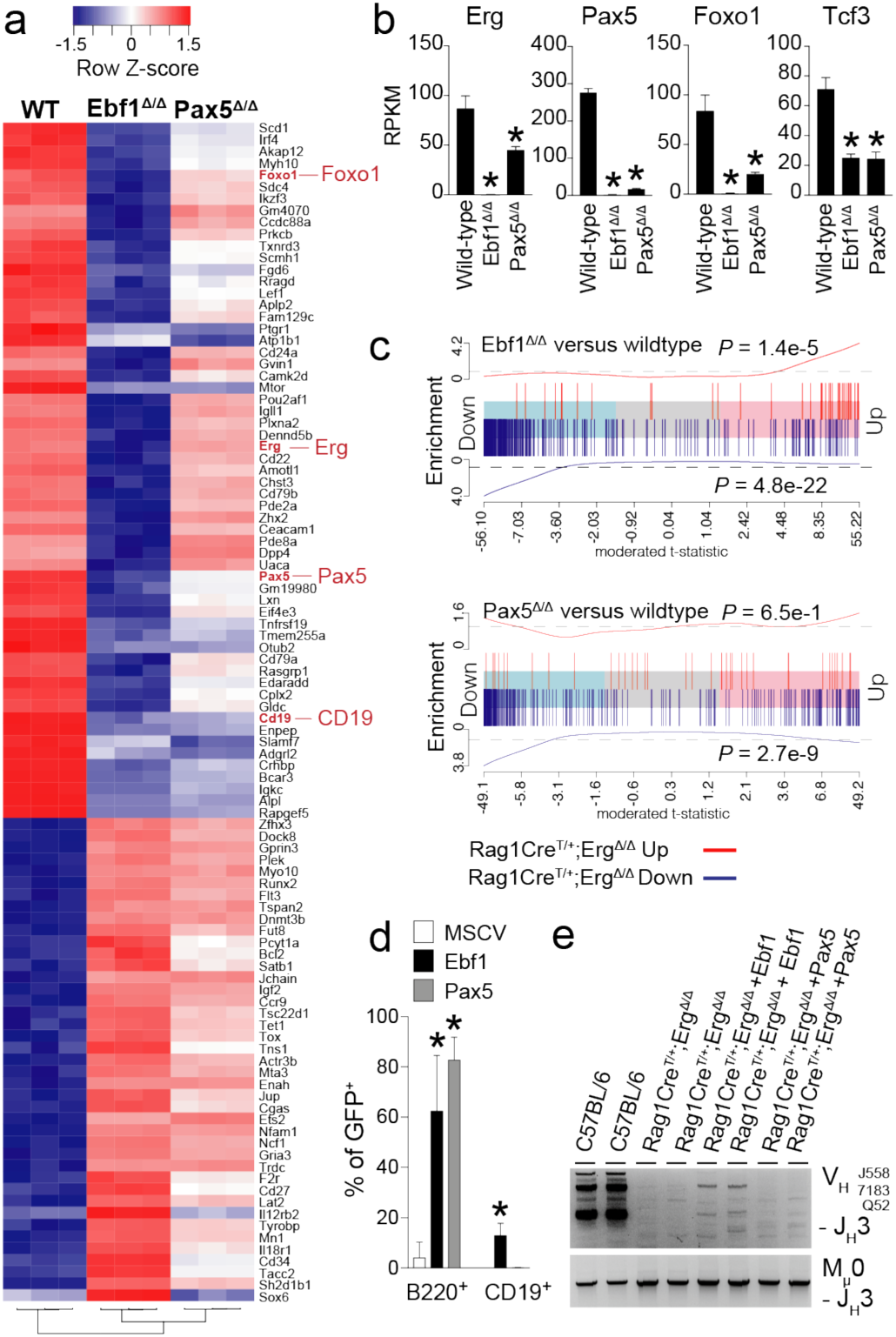
Gene expression in Ebf1- and Pax5-deficient B-cell progenitors and rescue of Erg-deficient B-cell progenitors. **A.** Heatmap of top 100 most variable genes in wild-type (n=3), Ebf^Δ/Δ^ (n=3), Pax5^Δ/Δ^ (n=3) B-cell progenitors with hierarchical clustering applied. **B.** Expression of *Erg*, *Pax5*, *Foxo1* and *Tcf3* in wild-type, Ebf^Δ/Δ^ and Pax5^Δ/Δ^ B-cell progenitors (RPKM). **C.** Barcode enrichment plots depicting strongly associated gene expression signatures of down (vertical blue bars) and up (vertical red bars) regulated genes in *Rag1Cre^T/+^;Erg^Δ/Δ^* pre-proB cells compared to Ebf1^Δ/Δ^ (top) and Pax5^Δ/Δ^ (bottom) B-cell progenitors. Genes are ordered (from left to right) as most downregulated to most upregulated in Ebf1^Δ/Δ^ or Pax5^Δ/Δ^ B-cell progenitors compared to wild-type. The x-axis shows the moderated t-statistic in Ebf1^Δ/Δ^ or Pax5^Δ/Δ^ versus wild-type cells. A camera gene set test ^45^ confirms the correlation between *Rag1Cre^T/+^;Erg^Δ/Δ^* pre-proB cell and Ebf1^Δ/Δ^ or Pax5^Δ/Δ^ B-cell progenitor expression signatures with *P* values as shown for up- and down-regulated genes. **D.** Percentage of B220^+^ and CD19^+^ expressing GFP^+^ B-cell progenitors derived from lineage negative *Rag1Cre^T/+^;Erg^Δ/Δ^* BM transfected with MSCV control (n=3), Ebf1-(n=3) or Pax5-expressing (n=3) retroviruses and cultured on OP9 stromal cells with IL-7, SCF and Flt-ligand. *, *P* < 0.005 by Student’s unpaired t-test compared to MSCV control. **E.** V_H_-to-DJ_H_ recombination of V_H_558, V_H_7183, V_H_Q52 segments (top panel) recombination with Mu0 loading controls (bottom panel) in B220^+^ enriched B-cell progenitors derived from lineage negative C57BL/6 (n=2), *Rag1Cre^T/+^;Erg^Δ/Δ^* (n=2), and *Rag1Cre^T/+^;Erg^Δ/Δ^* BM transfected with Ebf1 (n=2) and Pax5 (n=2) retroviruses.

Finally, to confirm that *Ebf1* and *Pax5* were transcriptional regulators down-stream of Erg in pre-proB cells, transfection of *Rag1Cre*^T/+^*;Erg*^Δ/Δ^ progenitor cells with MSCV-driven constructs for constitutive expression of *Ebf1* and *Pax5* was performed. This experiment demonstrated rescue of B220 expression with *Ebf1* or *Pax5* over-expression in Erg deficient progenitors (Figure 5D). Notably, only partial rescue of CD19 expression and V_H_-to-DJ_H_ recombination was observed with *Ebf1* over-expression while no rescue was observed with *Pax5* over-expression (Figure 5D,E). These observations suggest that while Ebf1 over-expression could partially compensate for several aspects of B-cell development in the absence of Erg, Pax5 over-expression alone could not. This is in keeping with a hierarchical model highlighting the importance of Erg as a key mediator of the network.

Taken together, these experiments demonstrated the existence of a co-dependent transcriptional network between Erg, Ebf1 and Pax5, that co-regulate critical target genes at the pre-proB cells stage of B-cell development.

To further the delineate the directly regulated target genes in an Erg-dependent Ebf1 and Pax5 transcriptional network, we undertook mapping of ChIP-seq binding of Erg, Ebf1 and Pax5 to differentially expressed genes at the pre-proB cell stage of development in *Rag1Cre*^T/+^*;Erg*^Δ/Δ^ cells. We identified that the majority of these target genes demonstrated direct combinatorial binding of Erg, Ebf1 and/or Pax5 to annotated promoter regions, gene body enhancer/putative enhancer regions or putative distal enhancer regions of these genes (Figure 6A). Detailed examination of several key target genes for which expression was completely dependent on Erg in pre-proB cells identified direct binding of Erg to the promoter and enhancer regions for several pre-BCR components, including *CD19*, *Igll1, Vpreb1* and *CD79a.* This occurred with co-ordinate binding of Ebf1 and Pax5 to the regulatory regions of these genes ^15^ (Figure 6B). In addition, indirect regulation by Erg at the *Rag1/Rag2* locus was also identified, with down-regulation of expression of transcription factors that bind and regulate the *Rag2* promoter such *Pax5*, *Lef1* and *c-Myb* in *Rag1Cre*^T/+^*;Erg*^Δ/Δ^ pre-proB cells (Figure 4B) ^39^, as well as direct binding of Erg to the conserved B-cell specific *Erag* enhancer ^40^ (***Figure S4C***). Importantly, the loss of *Rag1* and *Rag2* expression in *Rag1Cre*^T/+^*;Erg*^Δ/Δ^ pre-proB cells occurred while expression of *Foxo1*, a positive regulator of the locus ^41^ was relatively maintained (Figure S4B).

**Figure 6.**
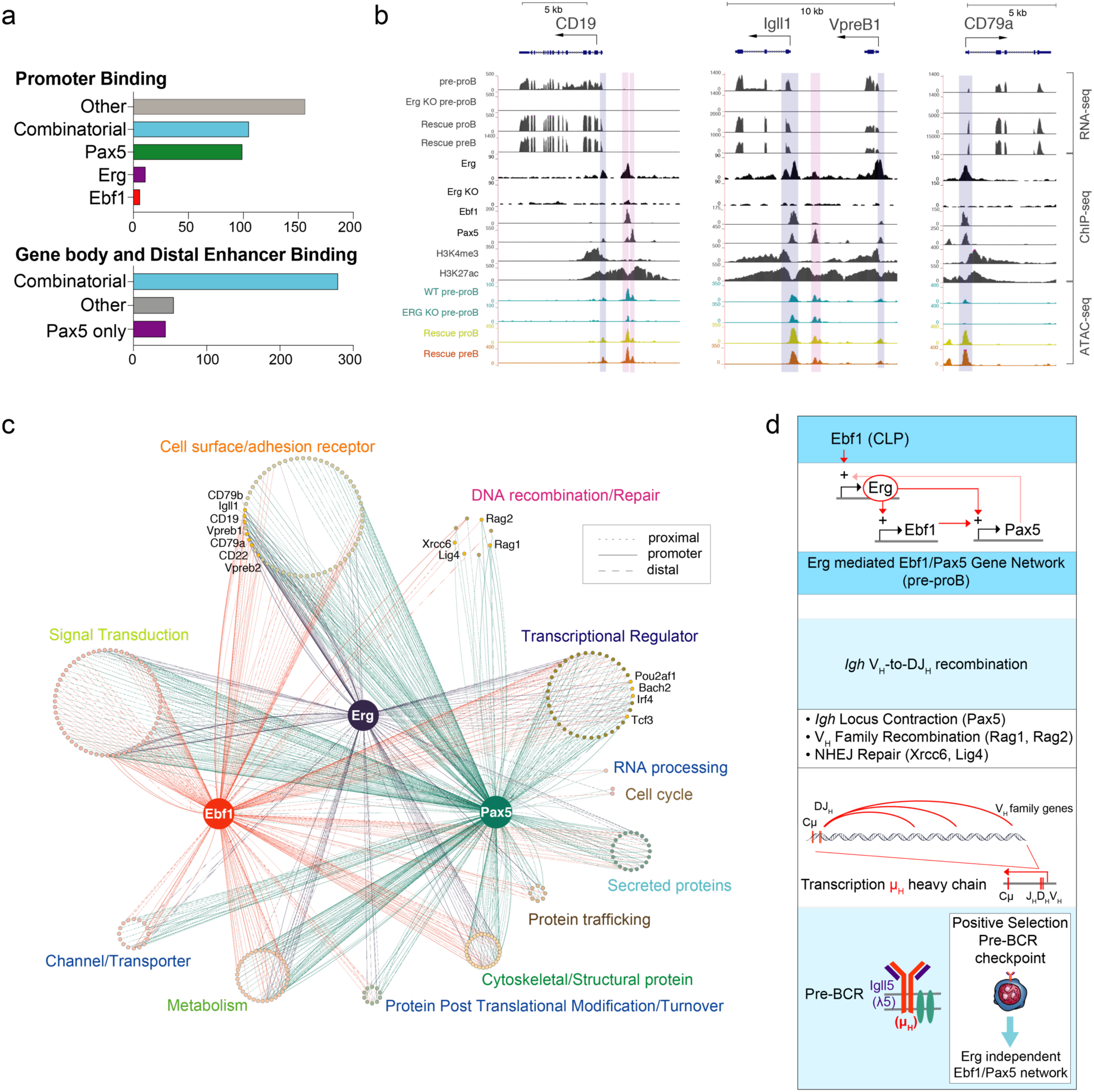
The Erg mediated Ebf1 and Pax5 gene regulatory network in pre-proB cells. **A.** Erg, Ebf1 and Pax5 binding to annotated regions of differentially expressed genes in *Rag1Cre^T/+^;Erg^Δ/Δ^* pre-proB cells. **B.** RNA-seq gene expression at *CD19, Igll1, VpreB1* and *CD79a* loci, with ChIP-seq of Erg, Ebf1, and Pax5 binding, H3K4me3 promoter mark, H3K27ac promoter and enhancer mark, and ATAC-seq of *Erg^fl/fl^* pre-proB cells, *Rag1Cre^T/+^;Erg^Δ/Δ^* pre-proB (Erg KO pre-proB), and Erg deficient proB and preB cells in *Rag1Cre^T/+^;Erg^Δ/Δ^;IgH^VH10tar/+^*mice rescued with a functionally rearranged immunoglobulin heavy chain allele. Solid blue bar: Erg, Ebf1 and/or Pax5 binding promoter. Solid pink bar: Erg, Ebf1 and/or Pax5 binding to enhancer regions. Open blue bar: promoter region with no binding of Erg, Ebf1 or Pax5. **C.** The Erg dependent Ebf1 and Pax5 transcriptional network in pre-proB cells with binding of each transcription factor shown to annotated promoter, proximal and distal gene regions of differentially expressed genes in *Rag1Cre^T/+^;Erg^Δ/Δ^* pre-proB cells. See ***Table S5***. **D.** Summary of the Erg dependent Ebf1 and Pax5 transcriptional network in V_H_-to-DJ_H_ recombination and pre-BCR formation.

An Erg-Ebf1-Pax5 mediated gene regulatory network was then mapped using each target gene, expression of which was perturbed in *Rag1Cre*^T/+^*;Erg*^Δ/Δ^ pre-proB cells, and that was directly bound by Erg, Ebf1 and/or Pax5 at promoter, proximal or distal gene regions, to provide a comprehensive representation of this gene network (Figure 6C).

A key observation arising from our data was that the B-cell developmental block arising in *Rag1Cre*^T/+^*;Erg*^Δ/Δ^ pre-proB cells could be overcome with the provision of a rearranged functional IgH VH10tar allele. This suggested that once the pre-BCR checkpoint was bypassed, *Erg* was no longer critical for further B-cell development and function, including V_L_J_L_ recombination of the *Igl* and BCR formation (Figure 3C,D). Indeed, beyond the pre-BCR checkpoint, re-emergence of *Ebf1* and *Pax5* expression occurred (Figure 4C) as well as expression of target genes of the *Ebf1* and *Pax5* network (Figure 6B, Figure S4B,C) in Erg-deficient *Rag1Cre*^T/+^*;Erg* ^Δ/Δ^;*IgH*^VH10tar/+^ proB and preB cells rescued with a VH10tar allele. This defines the role of Erg as an exquisitely stage specific regulator of early B cell development.

## Discussion

In this study we explored the role of the transcription factor *Erg* in B-lymphopoiesis. Our studies suggest two regions controlling Erg expression during B-cell development: the *Erg* promoter region and the H3K27ac-marked putative enhancer region in the first intron, to which the B-cell transcription factors Ebf1 and Pax5 directly bind. Complete loss of *Erg* expression in Ebf1^Δ/Δ^ Β-cell progenitors in which *Pax5* and *Foxo1* expression was also lost, place the initiation of Erg expression in the B-lymphoid lineage downstream of the E2A, Ebf1, Foxo1 transcriptional network at the CLP stage of lymphoid development ^10^. The importance of Erg in B-cell development was demonstrated in mice in which *Erg* had been deleted throughout lymphopoiesis, which exhibited a developmental block at the pre-proB cell stage that was associated with profound defects in V_H_-to-DJ_H_ recombination, *Igh* locus organization and transcriptional changes in multiple B-cell genes, including loss of expression of *Ebf1*, and *Pax5*. Combining RNA-seq, ChIP-seq and gene complementation studies, we were able to define a co-dependent transcriptional network between Erg, Ebf1 and Pax5, with direct Erg binding to the proximal (β) *Ebf1* promoter, to which Pax5, Ets1 and Pu.1 also co-operatively bind ^38^, as well as Erg binding to the *Pax5* promoter and potent intron 5 enhancer region, two critical *Pax5* regulatory elements required for correct transcriptional initiation of *Pax5* in early B-cell development ^12^. These data support a model (Figure 6D) in which Ebf1 expression, initially Erg-independent in CLPs, requires Erg in pre-proB cells to promote and maintain its expression. Erg is also required for simultaneous *Pax5* expression at this stage of development for the establishment of an inter-dependent B-lymphoid gene regulatory network.

Together Erg, Ebf1 and Pax5 directly co-regulated the expression multiple genes that had previously been identified as direct transcriptional targets of Ebf1 and Pax5 (Figure 6D). Direct Erg binding to promoters of the pre-BCR signalling complex genes such as *Igll1*, *VpreB* and *CD79a,* establish Erg as a transcriptional regulator of target genes in this network. In addition to *Rag1* and *Rag2*, we also identified network regulation of expression of *Xrcc6*, the gene encoding the Ku70 subunit of DNA-dependent protein kinase holoenzyme (DNA-PK) that binds DNA double strand breaks during V(D)J recombination ^42^, and *Lig4,* encoding the XRCC4 associated DNA-ligase that is required for DNA-end joining during V(D)J recombination ^43^ (Figure 6C, S4C). Along with direct Erg promotion of expression of *Pax5* as a structural regulator of the *Igh* locus, these findings are sufficient to explain the *Rag1Cre*^T/+^*;Erg*^Δ/Δ^ phenotype in which V_H_-toDJ_H_ recombination was lost. Together with loss of expression of components of the pre-BCR complex, we can conclude B-cell development was blocked as a consequence of *Erg* deletion due to the collapse of the Erg-mediated transcriptional network.

Importantly, re-emergence of *Ebf1* and *Pax5* expression beyond the pre-BCR checkpoint in *IgH*-rescued *Rag1Cre^T/+^;Erg^Δ/Δ^;IgH^VH10tar/+^* cells was observed, along with expression of target genes of Ebf1 and Pax5. This demonstrates that Erg is a stage-specific regulator of B-cell development, with emergence of an Erg-independent Ebf1 and Pax5 gene network during later stages of B-cell development, once clones have transitioned through the pre-BCR checkpoint. This would allow IgL chain V_L_ to J_L_ recombination and BCR formation to proceed in preB cells in which endogenous *Erg* expression is also reduced (Figure 1B,C). *Erg* however, is critical for promoting *Ebf1* and *Pax5* expression in pre-pro-B cells, orchestrating a transcriptional network required for V_H_-to-DJ_H_ recombination, pre-BCR formation, and early B cell development. In this role, Erg not only co-ordinates the transcriptional functions of Ebf1 and Pax5, but reinforces the *Erg*-mediated transcriptional network by directly binding and activating critical target genes required for transition through the pre-BCR checkpoint.

## Acknowledgements

We thank Janelle Lochland, Jason Corbin, Jasmine McManus, Melanie Salzone, Carolina Alvarado, Keti Stoev, Nicole Lynch and Shauna Ross for skilled assistance. We thank Professor Robert Brink for the V_H_10tar knock-in mouse line. This work was supported by Program Grants (1113577, 1016647, 1054618, 1054925), Project Grant (APN 1060179, 1122783), Fellowship (DMT 1060675, SLN 1155342, WSA 1058344, TMJ 1124081), C.R.B. Blackburn Scholarship (MSYL, Australian National Health and Medical Research Council jointly with Royal Australasian College of Physicians) and Independent Research Institutes Infrastructure Support Scheme Grant (361646) from the Australian National Health and Medical Research Council, the Australian Cancer Research Fund and Victorian State Government Operational Infrastructure Support. YCC was supported through Maddie Riewoldt’s Vision. The MAGEC laboratory was supported by the Australian Phenomics Network and the Australian Government through the National Collaborative Research Infrastructure Strategy Program.

## Author Contribution

Conceptualization, A.P.N., M.S.Y.L, A.J.K., T.M.J., M.A.D., R.S.A., K.R., D.M.T., G.K.S., M.J.D., S.L.N. and W.S.A.; Methodology, A.P.N., M.S.Y.L, T.M.J., T.B., M.A.D., R.S.A., K.R., D.M.T., G.K.S., M.J.D., S.L.N. and W.S.A.; Investigation, A.P.N., H.C., S.H., K.B., T.M.J., M.S.Y.L., C.C.B., O.G., Y.C.C., T.B., L.D., C.D.H., H.I., S.M., E.V., T.W., K.R., G.K.S., M.J.D.; Formal analysis, A.P.N., H.C., S.H., M.S.Y.L., O.G., C.C.B., Y.C.C. T.B., K.R., M.J.D., S.L.N; Writing – Original Draft, A.P.N.; Writing – Review & Editing, A.P.N., H.C., S.H., G.K.S., S.L.N., and W.S.A.; Funding Acquisition, A.P.N. and W.S.A.; Supervision, A.P.N., M.A.D., D.M.T., G.K.S., M.J.D., S.L.N. and W.S.A.

## COMPETING FINANCIAL INTERESTS

The authors declare that there are no competing financial interests.

## Figure Legends

**Figure 1.** Expression of the *Erg* locus and targeted disruption of *Erg* in lymphopoiesis.

**Figure 2.** The immunoglobulin heavy chain locus in *Rag1Cre^T/+^;Erg^Δ/Δ^* mice.

**Figure 3.** A Rearranged *V_H_10_tar_* IgH allele rescues *Rag1Cre*^T/+^*;Erg^Δ/Δ^* B-lymphoid development.

**Figure 4.** Gene expression in *Rag1Cre^T/+^;Erg^Δ/Δ^* pre-proB cells and Erg DNA binding.

**Figure 5.** Gene expression in Ebf1- and Pax5-deficient B-cell progenitors and rescue of Erg-deficient B-cell progenitors.

**Figure 6.** The Erg mediated Ebf1 and Pax5 gene regulatory network in pre-proB cells.

## CONTACT FOR REAGENT AND RESOURCE SHARING

Further information and requests for resources and reagents should be directed to and will be fulfilled by the Lead Contact, Ashley Ng subject to Material Transfer Agreements.

## EXPERIMENTAL MODEL AND SUBJECT DETAILS

### Mice

Mice carrying the *Erg*^tm1a(KOMP)wtsi^ knock-first reporter allele ^46^ (*Erg^KI^*, KOMP Knockout Mouse Project) were generated by gene targeting in ES cells. Mice with a conditional *Erg* knockout allele (*Erg^fl^*) from which the IRES-LacZ cassette was excised was generated by interbreeding *Erg^KI^* mice with *Flpe* transgenic mice ^47^. *Rag1Cre* mice ^48^, in which Cre recombinase is expressed during lymphopoiesis from the CLP stage ^22^, were interbred with *Erg^fl^* mice to generate mice lacking *Erg* in lymphopoiesis (*Rag1Cre*^T/+^*;Erg*^Δ/Δ^) and *Rag1Cre*^+/+^*;Erg*^fl/fl^ (*Erg^fl/fl^*) controls. Mice carrying the rearranged immunoglobulin heavy chain *IgH^VH10tar^* allele ^49^ were a gift from Professor Robert Brink. *Rag1^-/-^* mice were obtained from the Jackson Laboratory. The cEμ^Δ/Δ^ and μΑ^Δ/Δ^ mice were generated by the MAGEC laboratory (Walter and Eliza Hall Institute of Medical Research) as previously described ^50^ on a C57BL/6J background. To generate cEμ^Δ^ mice, 20 ng/μl of Cas9 mRNA, 10 ng/μl of sgRNA (GTTGAGGATTCAGCCGAAAC and ATGTTGAGTTGGAGTCAAGA) and 40 ng/μl of oligo donor (CAAGCTAAAATTAAAAGGTTGAACTCAATAAGTTAAAAGAGGACCTCTCCAGTTTCGGCTCAACTCAACATTGCTCAATTCATTTAAAAATATTTGAAACTTAATTTATTATTGTTAAAA) were injected into the cytoplasm of fertilized one-cell stage embryos. To generate μΑ^Δ^ mice, 20 ng/μl of Cas9 mRNA, 10 ng/μl of sgRNA (GAACACCTGCAGCAGCTGGC) and 40 ng/μl of oligo donor (GCTACAAGTTTACCTAGTGGTTTTATTTTCCCTTCCCCAAATAGCCTTGCCACATGACCTGCCAGCTGCTGCAGGTGTTCTGGTTCTGATCGGCCATCTTGACTCCAACTCAACATTGCT) were injected into the cytoplasm of fertilized one-cell stage embryos. Twenty-four hours later, two-cell stage embryos were transferred into the oviducts of pseudo-pregnant female mice. Viable offspring were genotyped by next-generation sequencing. Mice were analysed from 5 to 14 weeks of age. Male and female mice were used. Experimental procedures were approved by the Walter and Eliza Hall Institute of Medical Research Animal Ethics Committee.

### Primary cell culture

B-cell progenitors were obtained from bone marrow that was lineage depleted using biotinylated Ter119, Mac1, Gr1, CD3, CD4, and CD8 antibodies, anti-biotin microbeads and LS columns (Miltenyi Biotec) and cultured on OP9 stromal cells in Iscove’s Modified Dulbecco’s Medium (Gibco, Invitrogen) supplemented with 10% (v/v) foetal calf serum (Gibco, Invitrogen), 50μM β-mercaptoethanol as well as murine interleukin-7 (10ng/mL) at 37°C in 10% CO_2_ for 7 days. Splenic B-cells were purified by negative selection using a B-cell isolation kit (Miltenyi Biotec) as described ^51^ and purity was confirmed by flow cytometry prior to labelling with Cell Trace Violet (CTV; Life technologies) as per manufacturer instructions. Labelled cells were seeded at 5×10^4^ cells per well and cultured for 90 hours.

## METHOD DETAILS

### Haematology

Blood was collected into tubes containing EDTA (Sarstedt) and analysed on an Advia 2120 analyser (Bayer).

### Flow Cytometry

Single-cell suspensions from bone marrow, lymph node or spleen were prepared in balanced salt solution (BSS-CS: 0.15M NaCl, 4mM KCl, 2mM CaCl_2_, 1mM MgSO_4_, 1mM KH_2_PO_4_, 0.8mM K_2_HPO_4_, and 15mM HEPES supplemented with 2% [vol/vol] bovine calf serum). Analysis of blood was performed after erythrocyte lysis in buffered 156mM NH_4_Cl. Staining was performed using biotinylated or fluorochrome-conjugated antibodies specific for murine antigens Ter119 (Ly-76), CD41 (MWReg30), Gr1 (Ly6G and Ly6C), Mac1 (CD11b), NK1.1, CD11c (N418), CD45R/B220 (RA3-6B2), CD19 (1D3), CD3 (17A2), CD4 (GK1.5), CD8a (53.6.7), Sca1 (Ly6A/E, D7), cKit (CD117, ACK4 or 2B8), CD150 (TC15-12F12.2), CD105 (MJ7/18), CD16/32 (24G2), CD127 (A7R34), CD135 (A2F10), Ly6D (49-H4), CD21/CD35 (7G6), CD23 (B3B4), CD93 (AA4.1), CD24 (M1/69), CD43 (S7), CD45.2 (S450-15-2), CD45.1 (A20), IgM^b^ (AF6-78), IgD (11-26c.2a), CD138 (281.2), IgG1 (X56), CD25 (3C7), CD44 (IM7). Secondary staining used streptavidin PE-Texas-Red (Invitrogen). FACS-Gal analysis was performed using warm hypotonic loading of fluorescein di β-D-galactopyranoside (Molecular Probes) on single cells as described ^52^ followed by immunophenotyping using relevant surface antigens as defined in Table S1. Cells were analyzed using a LSR II or FACS Canto flow cytometer (Becton Dickinson) or sorted using a FACSAria II (Becton Dickinson) flow cytometer after antibody staining and lineage selection or depletion using anti-biotin beads and LS columns (Miltenyi Biotec). Data was analysed using FlowJo software (Version 8.8.7, Tree Star).

### Genomic PCR

Genomic DNA was extracted using DirectPCR lysis reagent (Viagen) with proteinase K (Sigma-Aldrich) or the DNeasy minikit (Qiagen). 1μL of supernatant from murine tail samples lysed in 200μL or 50-100ng of genomic DNA were used for each reaction. Conditional *Erg* genomic deletion was detected using primers designed to detect the wild-type, floxed or deleted *Erg* alleles (Table S3). Degenerate PCR primers for detection of genomic recombination across distal V_H_558 or proximal V_H_7183, V_H_Q52 regions, the D_H_ region or Mu0 regions to J3 segments ^53^, and Vκ ^54^ were used as described, as were primers to detect TCR VβJ recombination ^55, 56^ and the *IgH^VH10tar^* allele ^57^ (Table S3) ^58 59–61^. PCR products were separated by agarose gel electrophoresis and visualized with ethidium bromide staining. For quantitative genomic PCR using SYBR green (Life technologies), primers spanning individual *Erg* exons were used as described ^19^ and relative quantitation was performed using the 2^-ΔΔCT^ method ^62^.

### Splenic B-cell culture

Splenic B-cells were purified and purity was confirmed by flow cytometry prior to labelling with Cell Trace Violet (CTV; Life technologies) as per manufacturer instructions. Labelled cells were seeded at 5×10^4^ cells per well and cultured for 90 hours with either AffiniPure F(ab’)₂ Fragment Goat Anti-Mouse IgM µ Chain Specific (20mg/ml; Jackson Immunoresearch), CD40L (produced in-house as described ^63^) supplemented with IL4 (10ng/ml; R&D systems) and IL5 (5ng/ml; R&D systems) to assess T-cell dependent responses, or LPS (25mg/ml; Difco) to assess T-cell independent responses, and analysed by flow cytometry.

### RNA-seq of primary B-cell progenitor samples

Total RNA was extracted using the RNeasy Plus minikit (Qiagen) from bone marrow B-lymphoid populations sorted independently from two *Rag1Cre*^T/+^*;Erg^Δ/Δ^* and *Rag1Cre*^+/+^*;Erg*^fl/fl^ mice at 7-10 weeks of age. Sequencing was performed on an Illumina Hi-Seq 2500 to generate 100bp paired-end reads. Two biological replicates were sequenced for each mouse strain and B-cell development stage. Adapter sequences were removed using Trimgalore (https://github.com/FelixKrueger/TrimGalore). Reads were aligned to the mm10 mouse genome using STAR ^64^. Genewise counts were obtained using featureCounts ^65^ with Rsubread’s built-in Entrez Gene annotation ^66^. Downstream analysis as conducted using edgeR 3.22.5 ^67^. For each B-cell stage, genes were filtered as non-expressed if they were assigned 0.5 counts per million mapped reads (CPM) in fewer than two libraries. Library sizes were TMM normalized and differential expression was assessed using quasi-likelihood F-tests ^68^. Genes were called differentially expressed if they achieved a false discovery rate of 0.05 (***Table S4***). For plotting purposes, counts were converted to Fragments Per Kilobase of transcript per Million mapped reads (FPKM) using edgeR’s rpkm function. These data have been deposited in Gene Expression Omnibus database (accession number GSE132854).

### Analysis of publicly available RNA-seq datasets

FASTQ files containing RNA-seq profiles of B-cell progenitor cells were downloaded from GEO for Ebf1^Δ/Δ^ (GSM2879293, GSM2879294, GSM2879295), Pax5^Δ/Δ^ (GSM2879296, GSM2879297, GSM2879298) and wildtype mice (GSM2879299, GSM2879300, GSM2879301). Reads were aligned to the mm10 genome using Rsubread’s align function and read counts were summarized at the gene level as for the primary B-cell samples ^66^. Genes were filtered from downstream analysis using edgeR’s filterByExpr function and library sizes were TMM normalized. Counts were transformed to log2-CPM and the mean-variance relationship estimated using the *voom* function in limma ^69^. Heatmaps were generated using heatmap.2 function in gplots. Genes were tested for differential expression using linear modelling in limma 3.38.2 ^70^. Gene set testing was performed using *camera* ^45^ and barcode plots were generated with limma.

### Chromatin Immunoprecipitation (ChIP)

Chromatin immunoprecipitation was performed on 2×10^7^ cultured ProB cells or primary *Rag1Cre*^T/+^*;Erg^Δ/Δ^* thymocytes as a negative control for Erg binding. Cells were cross-linked with 1% formaldehyde for 15 min at room temperature, terminated by the addition of 0.125M glycine. Cells were then lysed in 1% SDS, 10mM EDTA, 50mM Tris-HCl, pH8.0, and protease inhibitors. Lysates were sonicated in a Covaris ultrasonicator to achieve a mean DNA fragment size of 500 bp. Immunoprecipitation was performed using 10μg of antibodies for a minimum of 12h at 4°C in modified RIPA buffer (1% Triton X-100, 0.1% deoxycholate, 90mM NaCl, 10mM Tris-HCl, pH8.0 and protease inhibitors). An equal volume of protein A and G magnetic beads (Life Technologies) were used to bind the antibody and associated chromatin. Reverse crosslinking of DNA was performed at 65°C overnight with RNaseA digestion followed by DNA purification using QIAquick PCR purification kits (Qiagen). Immunoprecipitated DNA was analysed on an Applied Biosystems StepOnePlus System with SYBR green reagents for iEμ μA and intergenic negative control regions using specific primers as detailed in Table S2. Relative ChIP PCR enrichment of the iEμ μA containing region in ProB cells compared to *Rag1Cre*^T/+^*;Erg^Δ/Δ^* thymocytes was performed and normalized to the intergenic negative control region using the 2^-ΔΔCT^ method ^62^.

### ChIP-seq

For sequencing analysis of immunoprecipitated DNA, DNA was quantified using the Qubit dsDNA HS Assay (Life Technologies). Library preparations were performed using the standard ThruPLEXTM-FD Prep Kit protocol (Rubicon Genomics) and size selected for 200–400bp fragments using Pippen Prep (Sage Science Inc.). Fragment sizes were confirmed using either the High Sensitivity DNA assay or the DNA 1000 kit and 2100 bioanalyzer (Agilent Technologies). Libraries were quantified with qPCR, normalized and pooled to 2nM before sequencing with single-end 75bp reads using standard protocols on the NextSeq (Illumina). DNA reads were adapter trimmed using Trimmomatic ^71^ and aligned to the GRCm38/mm10 build of the Mus musculus genome using the BWA aligner ^72^. Peaks were called using MACS2 ^73^ with default parameters to identify peaks using C17 antibody for Erg binding with *Rag1Cre*^T/+^*;Erg^Δ/Δ^* thymocytes as a negative control to filter peaks not due to Erg binding, and were annotated to closest (peak start within 10kb from TSS) and overlapping genes using the R/Bioconductor package ChIPpeakAnno ^74^ (***Table S5***). These data have been deposited in Gene Expression Omnibus database (accession number GSE132853). Publicly available FASTQ files for Ebf1 (GSM1296532, GSM1296537), Pax5 (GSM932924), H3K4me3 (GSM2255547) and H3K27Ac (GSM2255552) ChIP-seq experiments were aligned to the mm10 mouse reference genome (GRCm38, December 2011) using Rsubread ^75^. Peak-calling was performed using MACS2 ^73^ against input FASTQ files (GSM1296537, GSM1145867). Coordinates for Pu.1, Pax5, Irf4, YY1, Rad21 and CTCF binding were as published ^76^, while coordinates from annotated immunoglobulin heavy chains were obtained from Ensemble/Biomart (<u>accessed 6th March 2017)</u> and coordinates for 3’ regulatory region (3’RR) hypersensitivity regions, 3’αE, iEμ, were as published ^77 78 79 14^.

### ATAC-seq analysis

ATAC-seq ^80^ was performed on sorted pre-proB, proB and preB populations. Briefly, 5×10^4^ nuclei were fragmented by sonication for 30 minutes at 37°C and the DNA purified prior to amplification with indexing primers (HiFi Ready Mix, Kapa Biosciences) for 13 PCR cycles followed by quality assessment by Bioanalyser. High quality libraries were size selected (150 – 700 base pairs) and sequenced using a high output paired end 75 base pair kit on the Nextseq 500 (Illumina) to a minimum of 50 million reads. ATAC-seq reads were aligned to mm10 genome using Bowtie2 ^81^ (http://bowtie-bio.sourceforge.net/bowtie2/index.shtml accessed 6th March 2017). Peak calling was performed using MACS2 ^73^. Intersections of genetic coordinates were performed using Bedtools (http://bedtools.readthedocs.io/en/latest/ accessed 6^th^ March 2017). Heatmaps of unique peaks were generated using pHeatmap in R. These data have been deposited in Gene Expression Omnibus database (accession number GSE132852).

### Visualisation of RNA-seq, ChIP-seq and ATASeq data

RNA-seq, ChIP-seq and ATAC-seq files were converted to BigWig files using deepTools (version 2) ^82^ and uploaded to Cyverse (www.cyverse.org) for visualization in UCSC Genome Browser ^83^ (genome.ucsc.edu).

### Gene Network Analysis

All Ebf1, Pax5 and Erg ChIP-seq peaks mapping to differentially expressed genes in *Rag1Cre*^T/+^*;Erg^Δ/Δ^* pre-proB cells within 10Kb of the transcriptional start sites (TSS) were identified. Peaks inside the gene body were annotated as “proximal targets”, peaks overlapping the TSS were labelled as promoter regulated targets, peaks less than 3kb upstream or downstream of the TSS were labelled as putative promoter regulated targets, peaks more than 3kb upstream or downstream TSS were labelled as putative distal targets. Gene Ontogeny (GO) annotation of differentially expressed genes was performed and underwent expert manual curation. The network was constructed using ^84^ CRAN package, and exported to Cytoscape ^85^ for customization using RCy3 ^86^ R/Bioconductor package.

### Hi-C Analysis

*In situ* Hi-C was performed as previously described ^87^. The data preprocessing and analysis was performed as previously described with changes in parameters ^13^. In brief, primary immune cell libraries were generated in biological duplicates for each genotype. An Illumina NextSeq 500 was used to sequence libraries with 80bp paired-end reads to produce libraries of sizes between 42 million and 100 million valid read pairs. Each sample was aligned to the mm10 genome using the *diffHic* package v1.14.0 ^88^ which utilizes cutadapt v0.9.5 ^89^ and bowtie2 v2.2.5 ^81^ for alignment. The resultant BAM file was sorted by read name, the FixMateInformation command from the Picard suite v1.117 (https://broadinstitute.github.io/picard/) was applied, duplicate reads were marked and then re-sorted by name. Read pairs were determined to be dangling ends and removed if the pairs of inward-facing reads or outward-facing reads on the same chromosome were separated by less than 1000 bp for inward-facing reads and 6000 bp for outward-facing reads. Read pairs with fragment sizes above 1000 bp were removed. An estimate of alignment error was obtained by comparing the mapping location of the 3’ segment of each chimeric read with that of the 5’ segment of its mate. A mapping error was determined to be present if the two segments were not inward-facing and separated by less than 1000 bp, and around 1-2% were estimated to have errors. Differential interactions (DIs) between the three different groups were detected using the *diffHic* package ^88^. Read pairs were counted into 100 kbp bin pairs. Bins were discarded if found on sex chromosomes, contained a count of less than 10, contained blacklisted genomic regions as defined by ENCODE for mm10 ^90^ or were within a centromeric or telomeric region. Filtering of bin-pairs was performed using the filterDirect function, where bin pairs were only retained if they had average interaction intensities more than 5-fold higher than the background ligation frequency. The ligation frequency was estimated from the inter-chromosomal bin pairs from a 500 kbp bin-pair count matrix. The counts were normalized between libraries using a loess-based approach. Tests for DIs were performed using the quasi-likelihood (QL) framework ^68^ of the edgeR package. The design matrix was constructed using a one-way layout that specified the cell group to which each library belonged and the mouse sex. A mean-dependent trend was fitted to the negative binomial dispersions with the estimateDisp function. A generalized linear model (GLM) was fitted to the counts for each bin pair ^91^, and the QL dispersion was estimated from the GLM deviance with the glmQLFit function. The QL dispersions were then squeezed toward a second mean-dependent trend, using a robust empirical Bayes strategy ^92^. A p-value was computed against the null hypothesis for each bin pair using the QL F-test. P-values were adjusted for multiple testing using the Benjamini-Hochberg method. A DI was defined as a bin pair with a false discovery rate (FDR) below 5%.

DIs adjacent in the interaction space were aggregated into clusters using the diClusters function to produce clustered DIs. DIs were merged into a cluster if they overlapped in the interaction space, to a maximum cluster size of 1 Mbp. The significance threshold for each bin pair was defined such that the cluster-level FDR was controlled at 5%. Cluster statistics were computed using the *csaw* package v1.16.0 ^93^. Overlaps between unclustered bin pairs and genomic intervals were performed using the InteractionSet package ^94^. Plaid plots were constructed using the contact matrices and the plotHic function from the *Sushi R* package ^95^. The color palette was inferno from the *viridis* package ^96^ and the range of color intensities in each plot was scaled according to the library size of the sample. The plotBedpe function of the *Sushi* package was used to plot the unclustered DIs as arcs where the z-score shown on the vertical access was calculated as -log_10_(*p*-value). These data have been deposited in Gene Expression Omnibus database (accession number GSE133246).

### Fluorescence In Situ Hybridisation

Cultured B-cell progenitors were resuspended in hypotonic 0.075M KCl solution and warmed to 37°C for 20 minutes. Cells were pelleted and resuspended in 3:1 (vol/vol) methanol:glacial acetic acid fixative. Fixed cells were dropped onto coated Shandon^TM^ polysine slides (ThermoFisher Scientific) and air dried. The cells were hybridized with FISH probes (Creative Bioarray) at 37°C for 16 hours beneath a coverslip sealed with Fixogum (Marabu) after denaturation at 73°C for 5 minutes. Cells were washed at 73°C in 0.4x SSC/0.3%NP_40_ for 2 minutes followed by 2x SSC/0.1%NP_40_ for less than 1 minute at room temperature and air dried in the dark and cover slipped. Images of nuclei were captured on an inverted Zeiss LSM 880 confocal using a 63x/1.4 NA oil immersion objective. Z-stacks of images were then captured using the lambda scan mode, a 405 and a multi-band pass beam splitter (488/561/633). The following laser lines were used: 405, 488, 561 and 633 nm. Spectral data was captured at 8 nm intervals. In all cases, images were set up with a pixel size of 70 nm and an interval of 150 nm for z-stacks. Single dye controls using the same configuration were captured and spectra imported for spectral unmixing using the Zen software (Zen 2.3, Zeiss Microscopy). Unmixed data was then deconvolved using the batch express tool in Huygens professional software (Scientific Volume Imaging). Images were analysed using TANGO software ^97^ after linear deconvolution. Nuclear boundaries were extracted in TANGO using the background nuclear signal in the Aqua channel. A 3D median filter was applied and the 3D image projected with maximum 2D image projection for nuclei detection using the Triangle method for automated thresholding in ImageJ ^98^. Binary image holes were filled and a 2D procedure implemented to separate touching nuclei using ImageJ 2D watershed implementation. The 2D boundaries of the detected nuclei were expanded in 3D and inside each 3D delimited region, Triangle thresholding was applied to detect the nuclear boundary in the 3D space. Acquired images from immunofluorescent probes were first filtered using 3D median and 3D tophat filter to enhance spot-like structures followed by application of the “spotSegmenter” TANGO plugin with only the best 4 spots having the brightest intensity kept for analysis. The spots identified by TANGO were manually verified against the original immunofluorescent image to identify and record the correct distance computed by TANGO between the aqua and 5-Rox immunofluorescent probes for both *Igh* alleles within a nucleus.

### Statistical Analysis

Student’s unpaired two-tailed t-tests were used using GraphPad Prism (GraphPad Software), unless otherwise specified. Unless otherwise stated, a *P* value of < 0.05 was considered significant.

## KEY RESOURCES TABLE

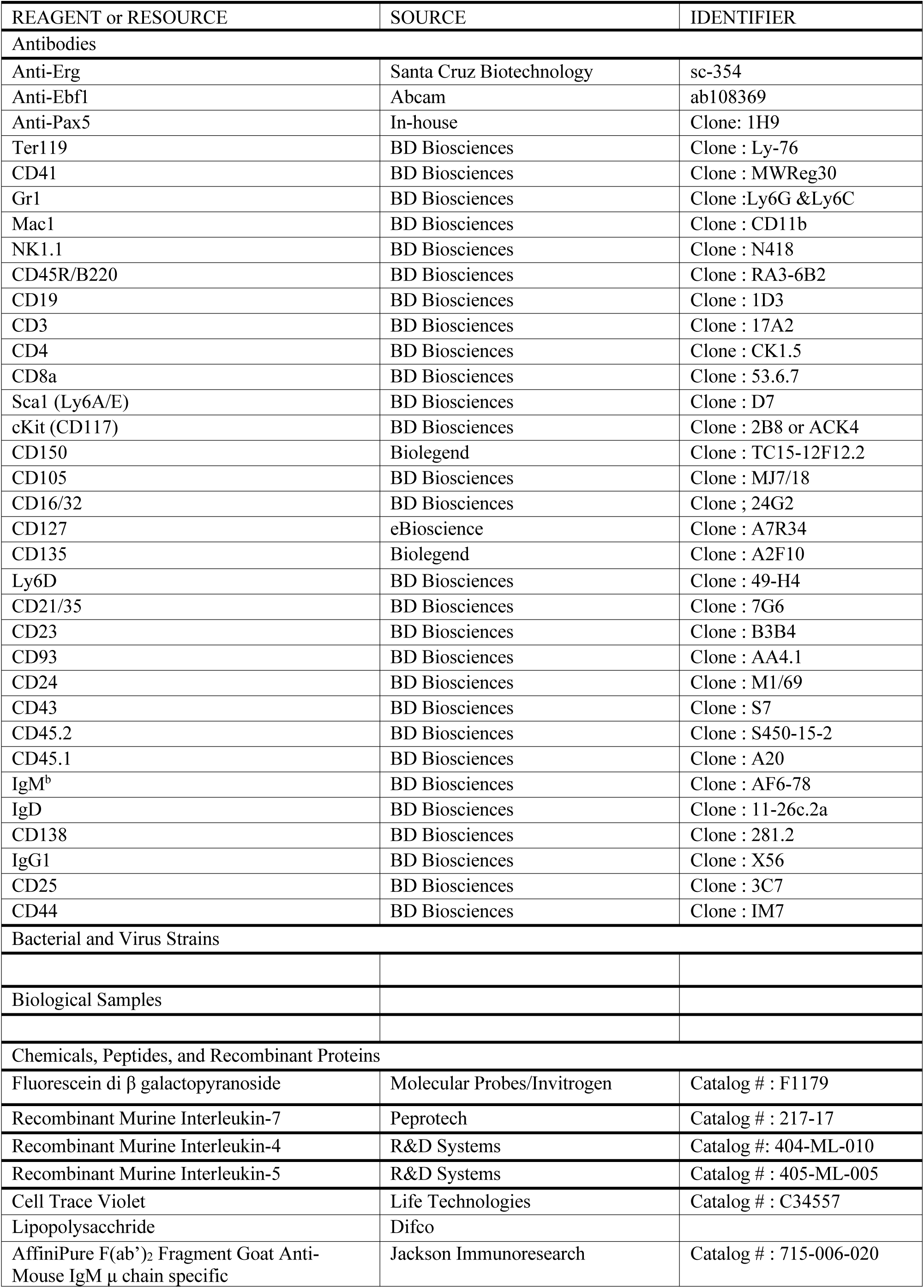
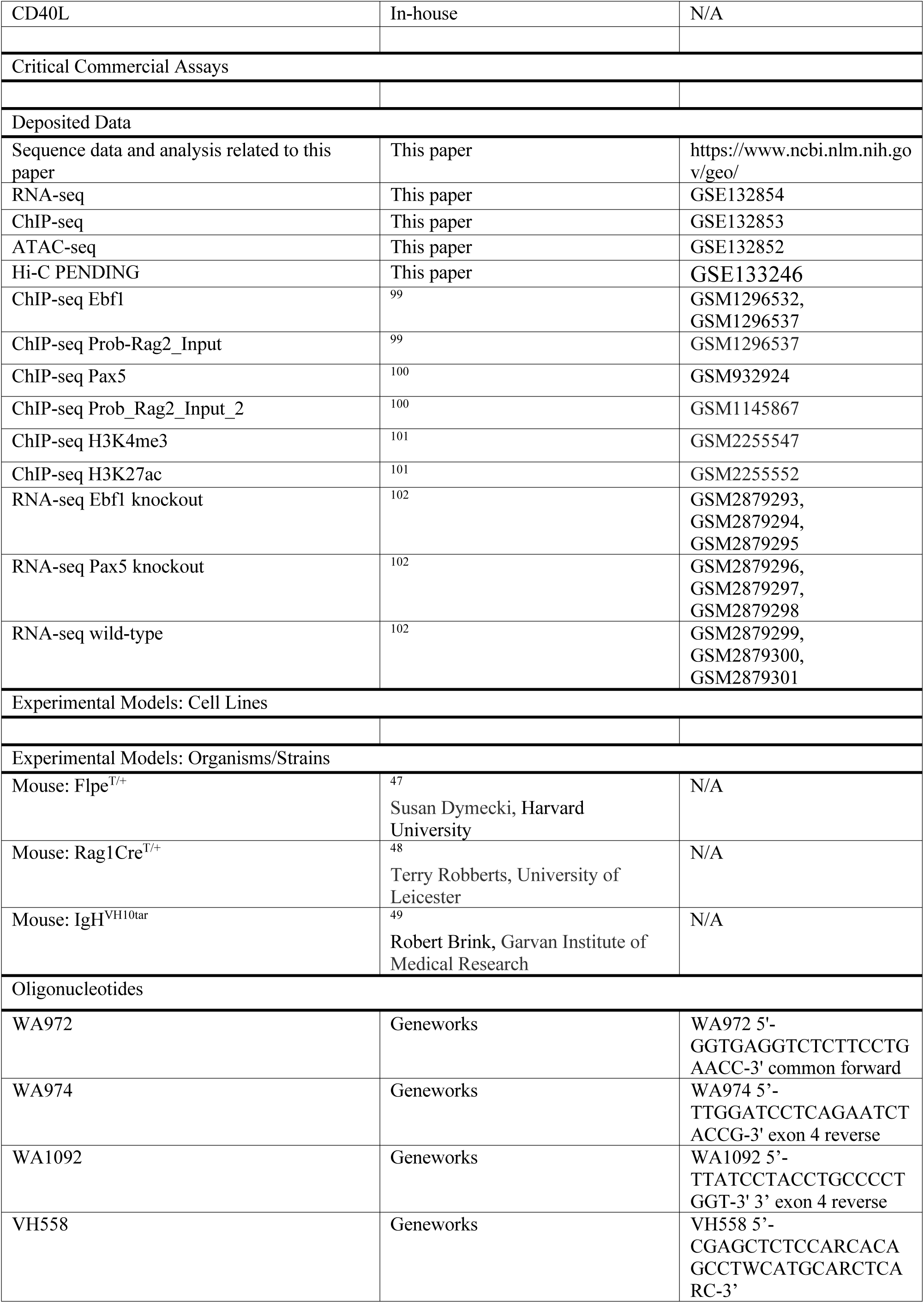
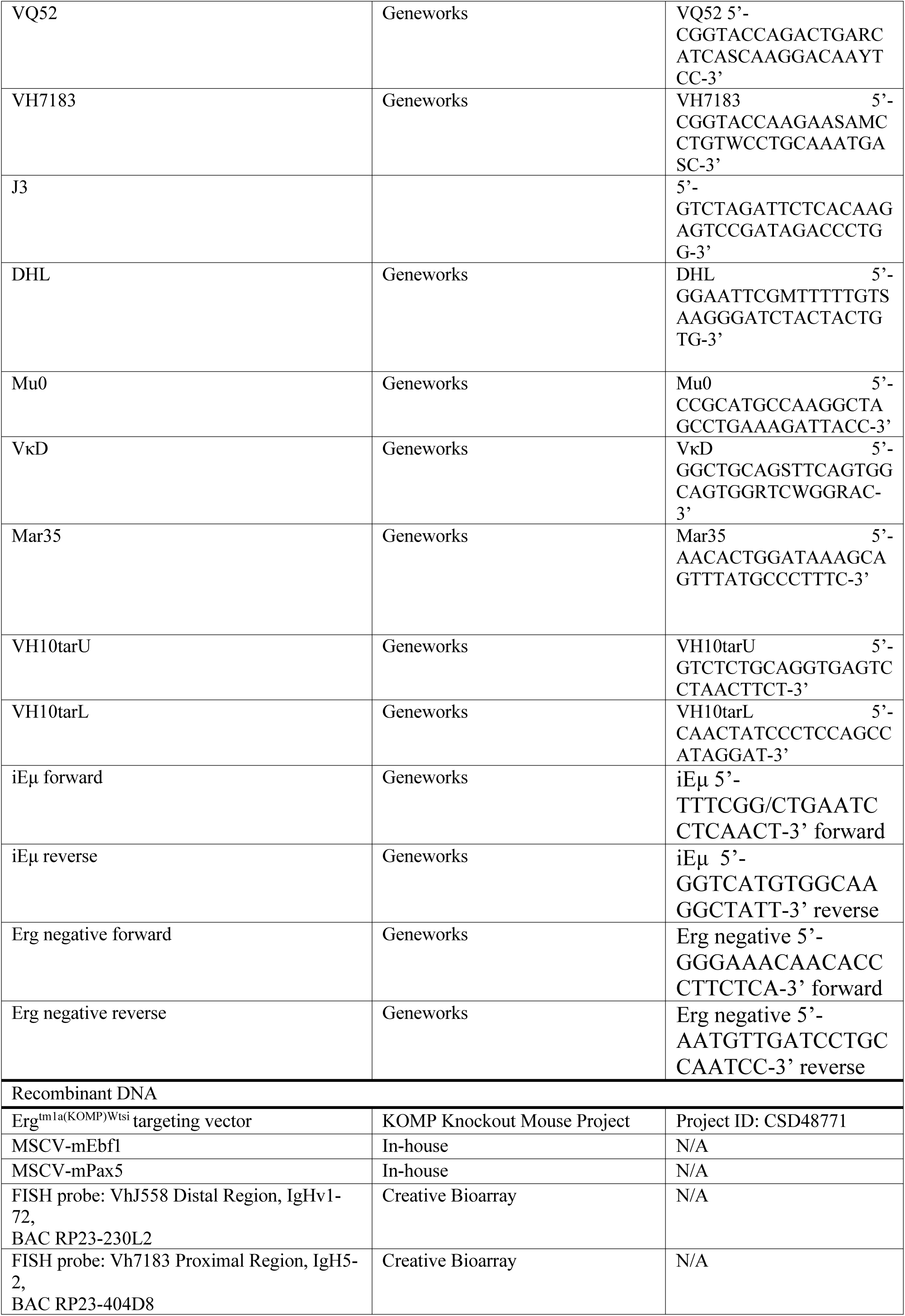
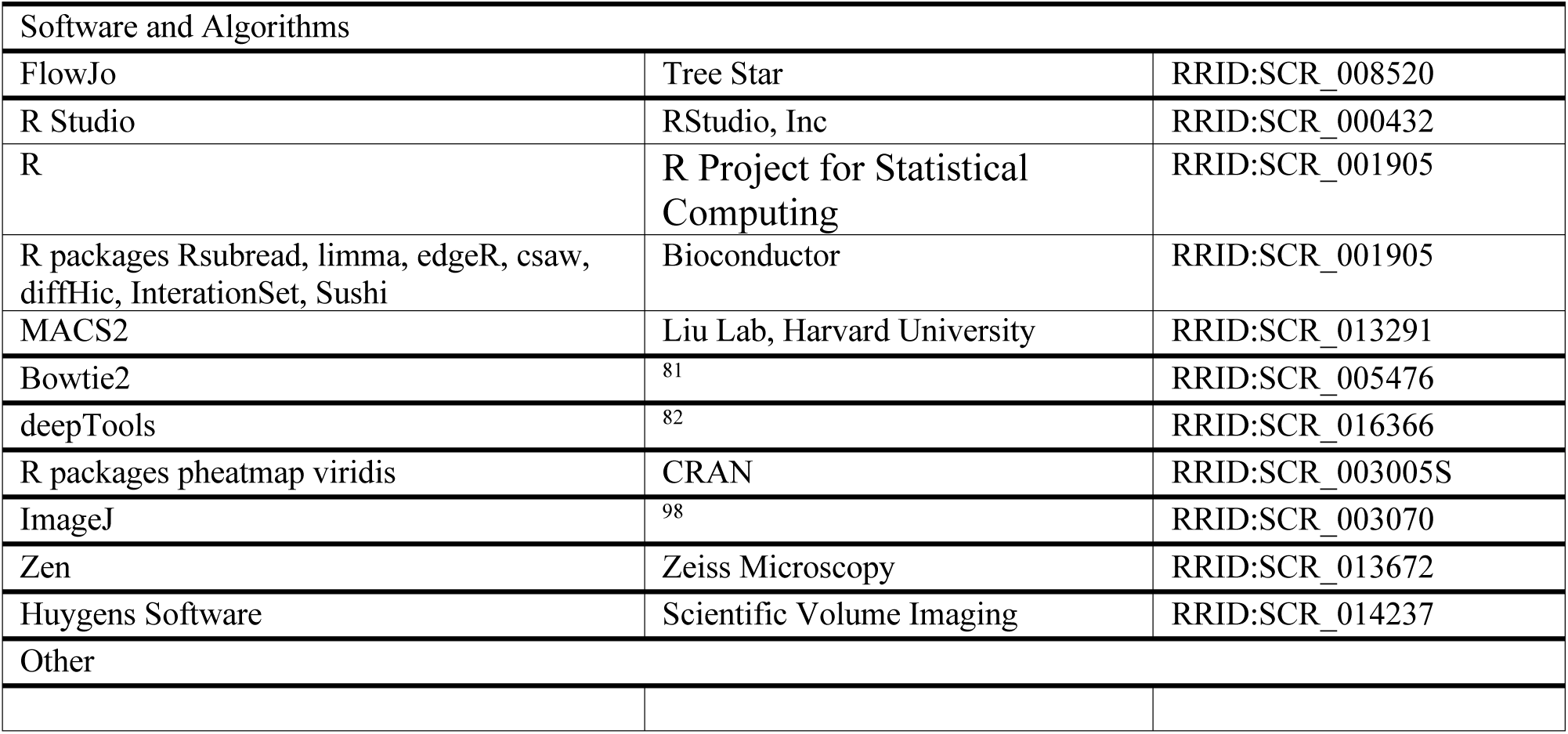

## Inventory of Supplemental Information

### Supplemental Figures

**Figure S1.** Representative flow cytometry plots indicating gating strategies for analysis of hematopoietic cell populations. Related to Figure 1

**Figure S2.** B-lymphopoiesis in *Rag1Cre^T/+^* mice and T-lymphopoiesis in *Rag1Cre^T/+^;Erg^Δ/Δ^* mice. Related to Figure 1.

**Figure S3.** In vivo Erg binding to the iEμ enhancer in pre-proB cells. Related to Figure 2.

**Figure S4.** RNA-seq and Erg, Ebf1 and Pax5 binding and chromatin accessibility at selected gene loci. Related to Figure 2, 4, 6.

### Supplemental Tables

**Table S1.** Immunophenotype of hematopoietic cell populations. Related to Figure 1.

**Table S2.** Peripheral blood counts of *Rag1Cre^T/+^;Erg^Δ/Δ^* mice. Related to Figure 1.

**Table S3.** Primers and PCR reactions. Related to Figure 2, 3.

**Table S4.** RNA-seq. Differentially expressed genes in *Rag1Cre^T/+;^Erg^Δ/Δ^* pre-proB cells, and Ebf1^Δ/Δ^ Pax5^Δ/Δ^ B-cell progenitors (EXCEL FILE). Related to Figure 4.

**Table S5.** Erg, Ebf1 and Pax5 ChIP binding coordinates to differentially expressed genes in *Rag1Cre^T/+;^Erg^Δ/Δ^* pre-proB cells (EXCEL FILE). Related to Figure 6.

---

**Figure S1.**
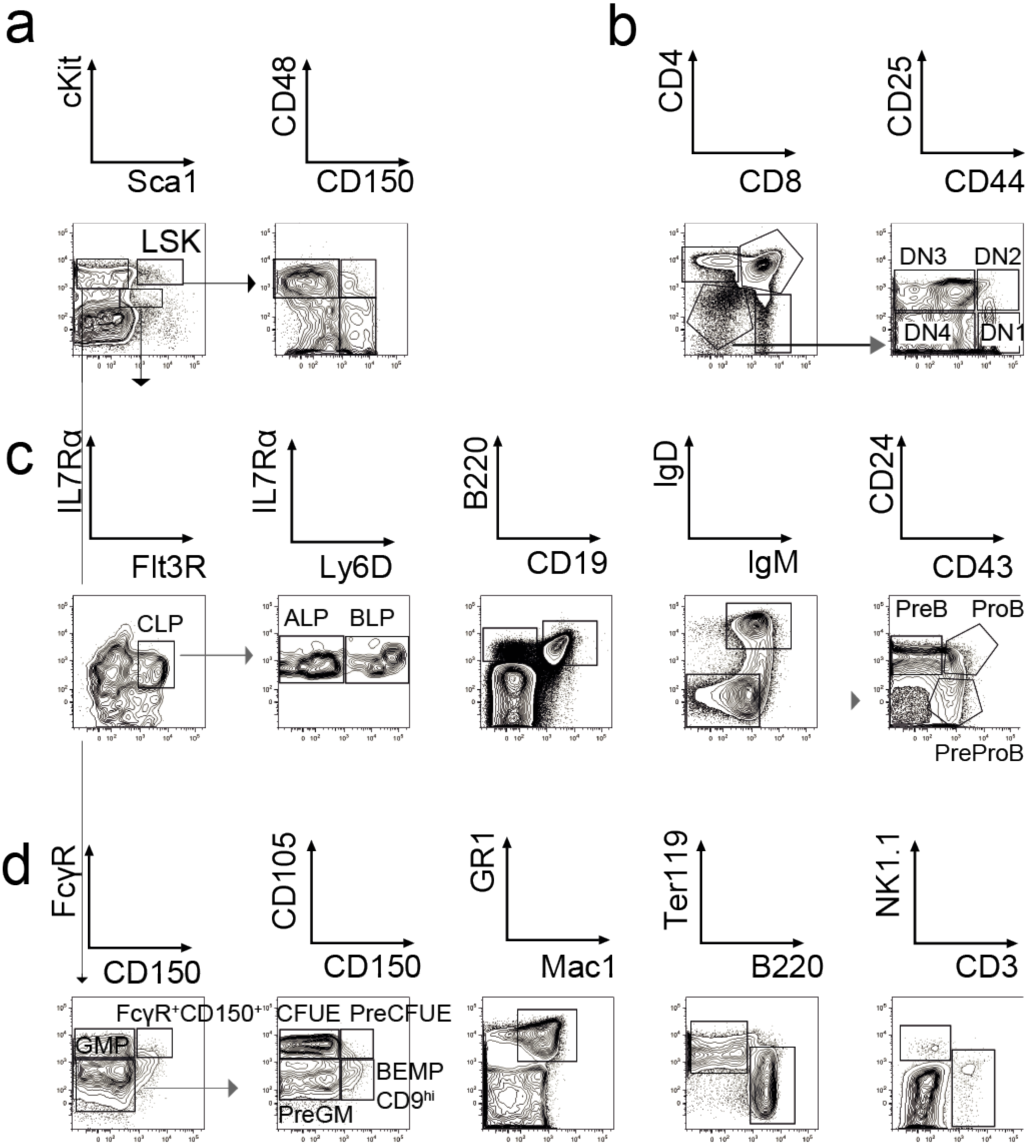
Representative flow cytometry plots indicating gating strategies for analysis of hematopoietic cell populations. **A**. bone marrow LSK cells, **B.** thymus sub-populations, **C.** bone marrow B-lineage cells and **D.** bone marrow myeloid cell populations in *Erg^KI/+^* mice. The cell surface markers and definitions of cell populations used are provided in Table S1.

**Figure S2.**
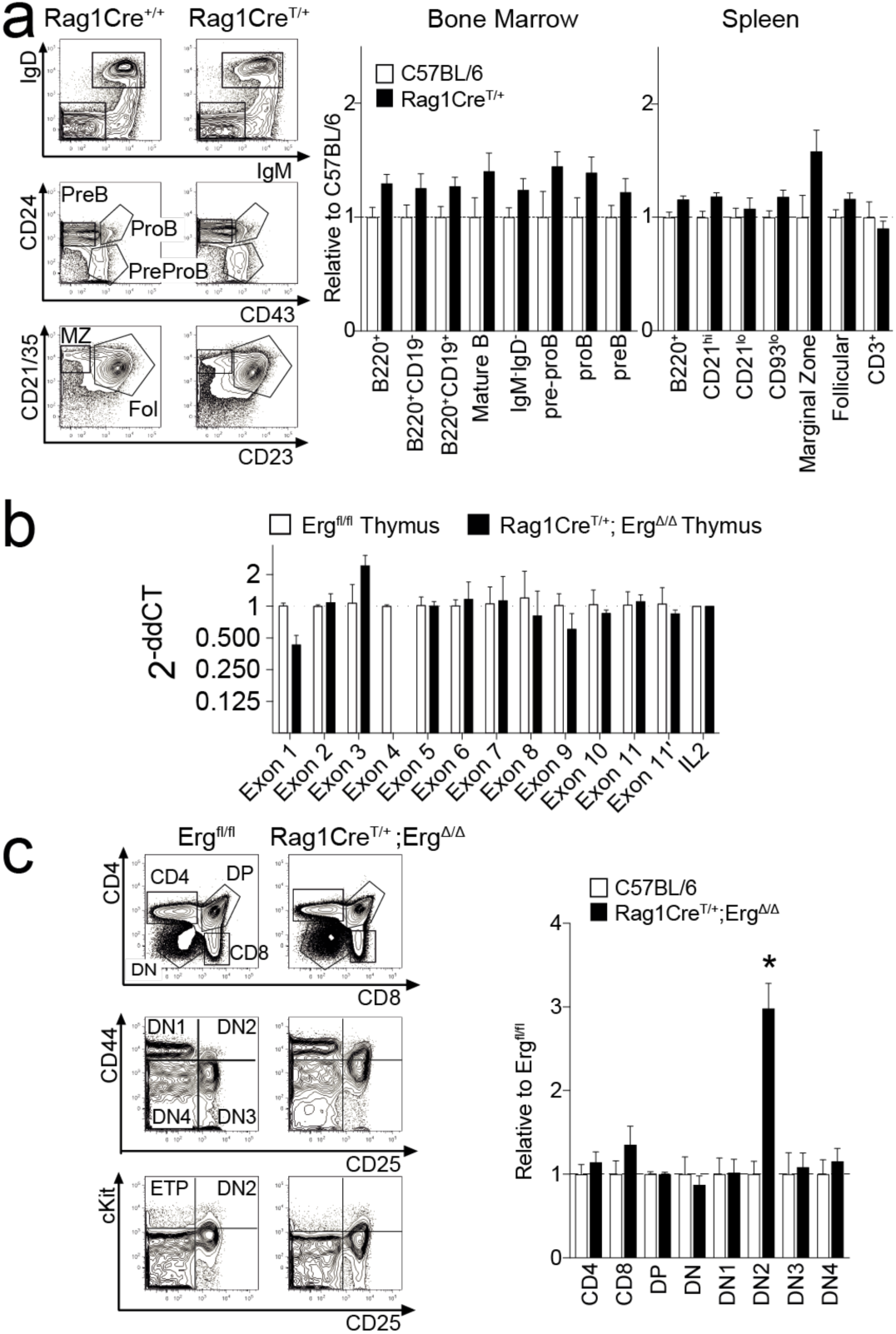
B-lymphopoiesis in *Rag1Cre^T/+^* mice and T-lymphopoiesis in *Rag1Cre^T/+^;Erg^Δ/Δ^* mice. **A.** Representative flow cytometry plots (left panels) of *Rag1Cre^T/+^* bone marrow and spleen cells. The IgM/IgD profile is from B220^+^ bone marrow cells (top panel), the CD24/CD43 profiles from B220^+^IgM^-^IgD^-^ bone marrow cells (middle panel) and the CD21/CD23 profiles from B220^+^CD93^lo^ spleen cells (bottom panel). Ratio of *Rag1Cre^+/+^* and *Rag1Cre^T/+^* B-lymphoid cells shown relative to *Rag1Cre^+/+^* controls in bone marrow and spleen with standard error of means (right panels). No significant differences were observed between the two genotypes for any population by Student’s two-tailed unpaired t-test corrected using Holm’s modification for multiple testing (*P_adj_* > 0.19, n=6 mice per genotype). **B.** Quantitative genomic PCR on DNA from *Erg^fl/fl^* and *Rag1Cre^T/+^;Erg^Δ/Δ^* thymocytes using primers spanning individual *Erg* exons ^19^ demonstrating efficient exon 4 deletion in *Rag1Cre*^T/+^*;Erg^Δ/Δ^* thymocytes by 2^-ΔΔCT^ method normalised to IL2 receptor and Erg exon 1. **C**. Representative flow cytometry plots (left panels) from *Erg^fl/fl^* and *Rag1Cre*^T/+^*;Erg^Δ/Δ^* thymi identifying the specific cell populations indicated with the mean with standard error of the mean of *Erg^fl/fl^* (n=7) and *Rag1Cre^T/+^;Erg^Δ/Δ^* mice (n=8) shown relative to the mean of *Erg^fl/fl^* controls (right panel). No significant differences were observed other that in the DN2 population (*P_adj_* < 6.3e-4 by Student’s two-tailed unpaired t-test corrected using Holm’s modification for multiple testing).

**Figure S3.**
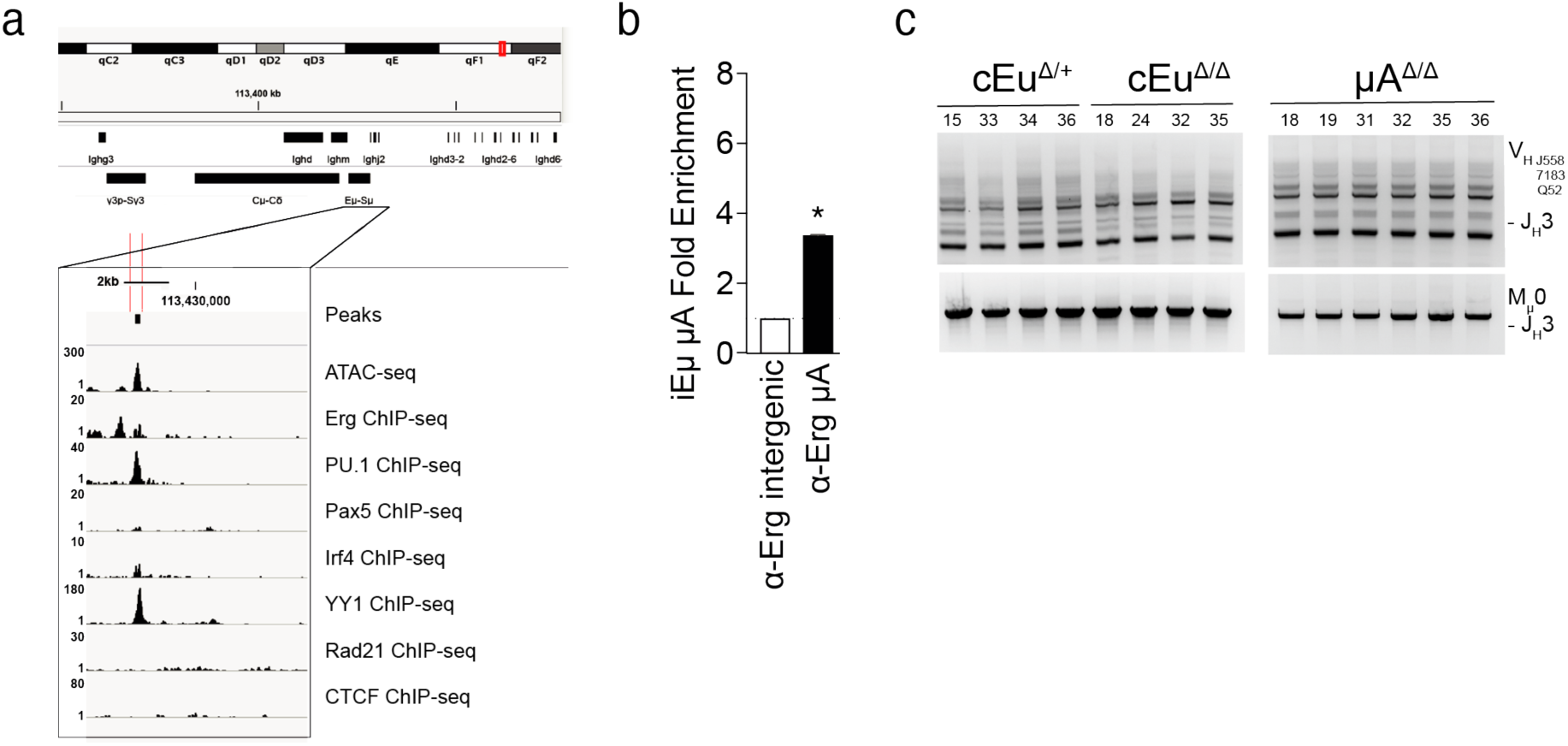
In vivo Erg binding to the iEμ enhancer. **A.** Region of the iEu enhancer as indicated (red bars) with the ATAC-seq and ChIP-seq for Erg, Pu.1, Pax5, Irf4, YY1, Rad21 and CTCF shown. **B.** Erg binding to iEμ containing the μA element by ChIP-PCR showing fold-enrichment in wild type B-cell progenitors relative to *Rag1Cre^T/+;^Erg^Δ/Δ^* thymocytes and normalised to a negative intergenic region control using the 2^-ΔΔCT^ method (n=2 biological replicates, * *P*<10^-4^ by Student’s two-tailed unpaired t-test). **C.** Genomic PCR using degenerate primers to IgH locus V_H_558, V_H_7183, V_H_Q52 segments for detection of V_H to_ DJ_H_ (top panel) and D_H_ to J_H_ (middle panel) recombination with Mu0 loading controls (bottom panel) in B220^+^ splenocytes.

**Figure S4.**
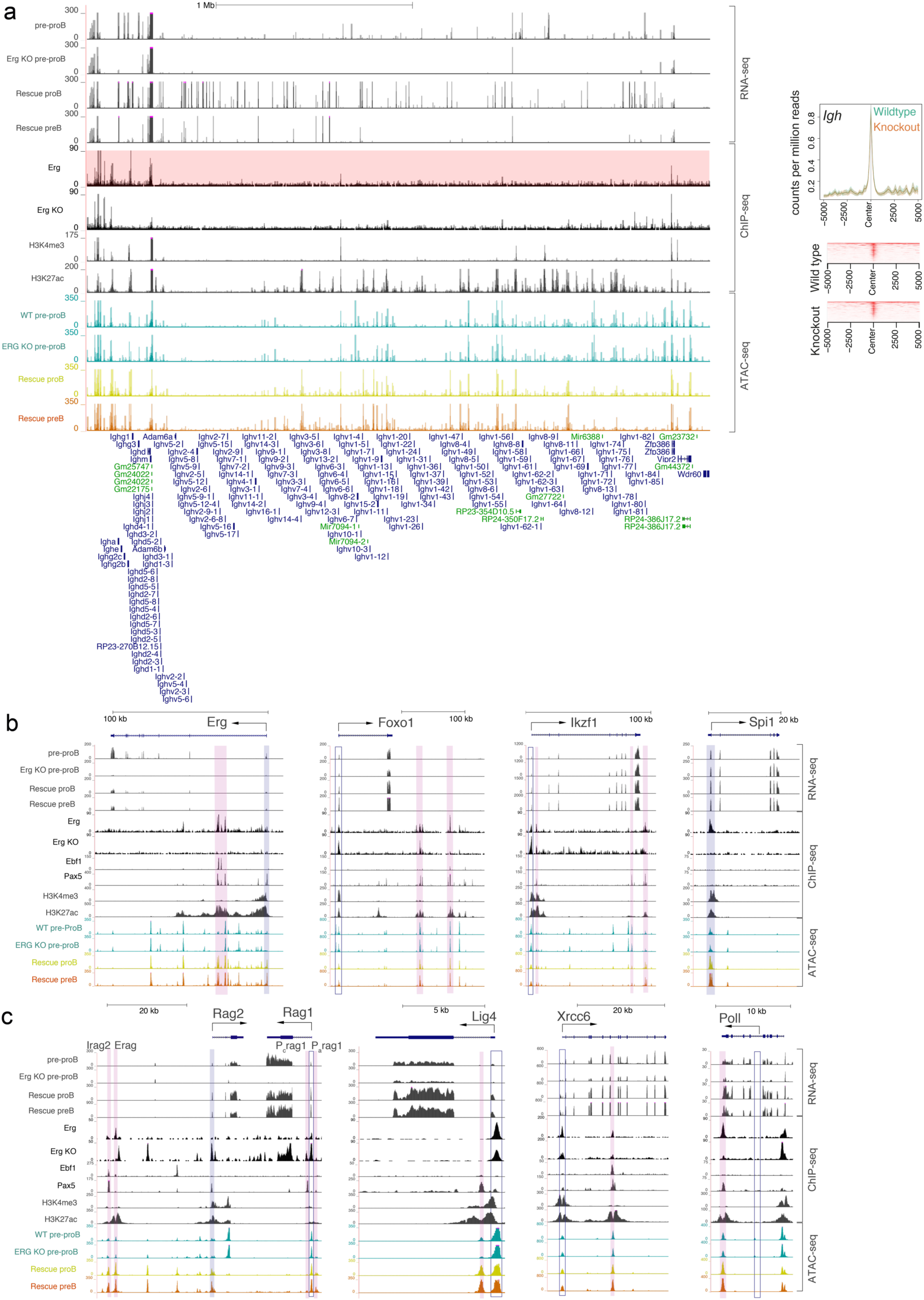
RNA-seq and Erg, Ebf1 and Pax5 binding and chromatin accessibility at selected gene loci. **A.** *Igh* locus with representative RNA-seq tracks shown for *Erg^fl/fl^* and *Rag1Cre^T/+^;Erg^Δ/Δ^* (Erg KO) pre-proB cells, and Erg deficient proB and preB cells in *Rag1Cre^T/+;^Erg^Δ/Δ^;IgH^VH10tar/+^*mice rescued with a functionally rearranged immunoglobulin heavy chain allele (Rescue proB, Rescue preB). ChIP-seq tracks for Erg (highlighted in pink), *Rag1Cre^T/+;^Erg^Δ/Δ^* thymus cells (Erg KO) to control for sites of non-Erg ChIP binding to DNA, H3K4me3 and H3k27ac shown. Chromatin accessibility by ATAC-seq (blue) in WT and Erg KO pre-proB cells, Erg deficient Rescue proB (yellow) and Rescue preB (orange) cells. **B and C.** RNA-seq gene expression at gene loci, with Erg, Ebf1, and Pax5 binding, H3K4me3 promoter mark, H3K27ac promoter and enhancer mark, and ATAC-seq in *Erg^fl/fl^* pre-proB cells (PreProB), *Rag1Cre^T/+;^Erg^Δ/Δ^* pre-proB (Erg KO pre-proB), and Erg deficient proB and preB cells in *Rag1Cre^T/+;^Erg^Δ/Δ^;IgH^VH10tar/+^*mice rescued with a functionally rearranged immunoglobulin heavy chain allele (Rescue proB, Rescue preB). Solid blue bar: Erg, Ebf1 and/or Pax5 binding promoter. Solid pink bar: Erg, Ebf1 and/or Pax5 binding to enhancer regions. Open blue bar: promoter region with no binding of Erg, Ebf1 or Pax5. **B.** *Erg, Foxo1, Ikzf1* and *Spi1* B-lineage transcription factor loci. **C.** *Rag1* and *Rag2*, *Lig4, Xrcc6* and *Poll* loci.

## Supplementary Tables

**Table S1.** Immunophenotype of hematopoietic cell populations. See Figure 1.

**Table S2.** Peripheral blood counts of *Rag1Cre^T/+^;Erg^Δ/Δ^* mice. See Figure 1.

**Table S3.** Primers and PCR reactions. See Figure 2, 3.

**Table S4.** RNA-seq. Differentially expressed genes in *Rag1Cre^T/+;^Erg^Δ/Δ^* pre-proB cells, and Ebf1^Δ/Δ^ Pax5^Δ/Δ^ B-cell progenitors (EXCEL FILE). See Figure 4.

**Table S5.** Erg, Ebf1 and Pax5 ChIP binding coordinates to differentially expressed genes in *Rag1Cre^T/+;^Erg^Δ/Δ^* pre-proB cells (EXCEL FILE). See Figure 6

**Supplementary Table S1.**
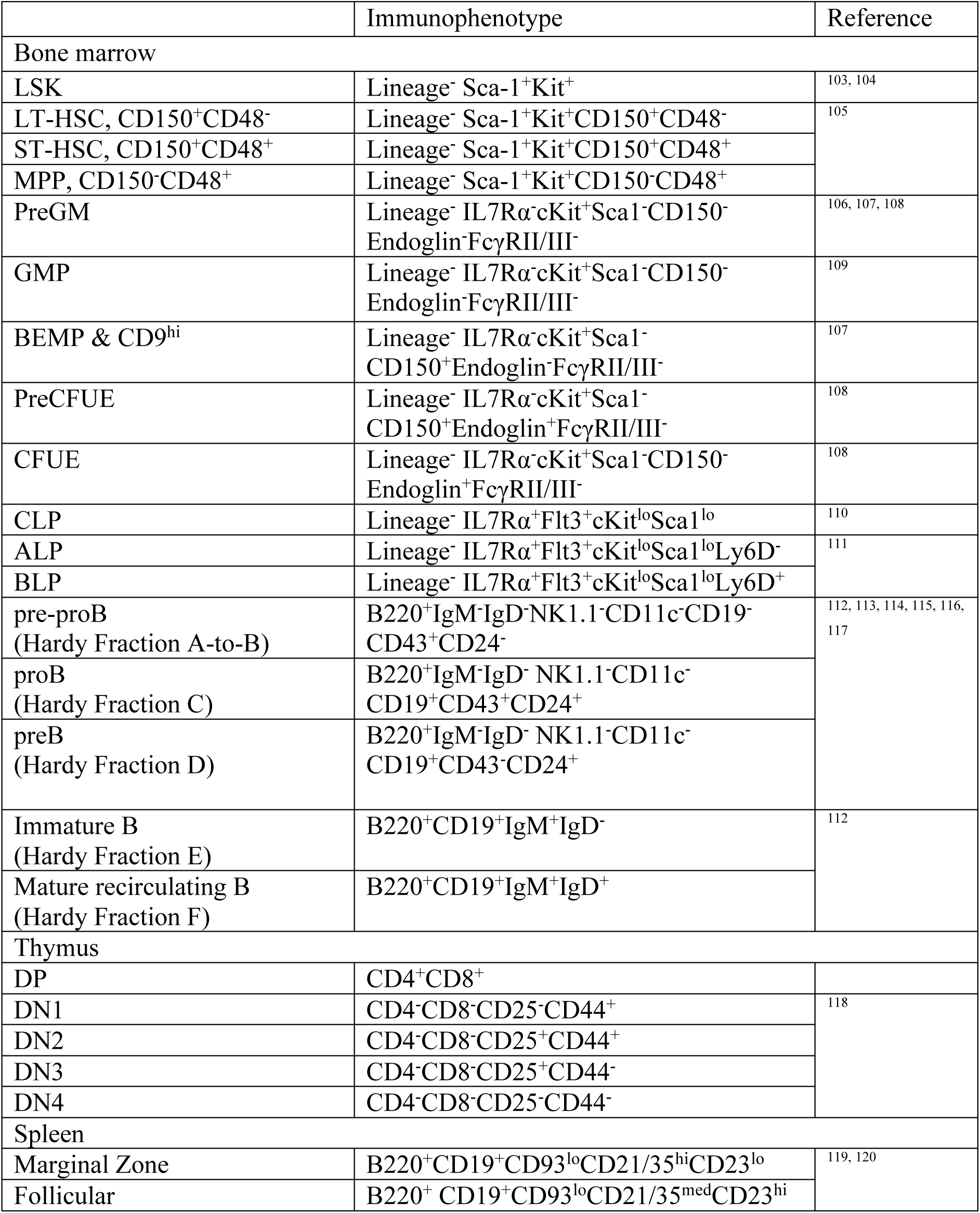
Immunophenotype of hematopoietic cell populations. See **Figure 1**

**Supplementary Table S2.** Peripheral blood counts of *Rag1Cre^T/+^;Erg^Δ/Δ^* mice. See **Figure 1**

| Genotype | RBC<br>x10<sup>12</sup>/L | Platelets<br>x10<sup>9</sup>/L | WBC<br>x10<sup>9</sup>/L | Neutrophil<br>x10<sup>9</sup>/L | Lymphocyte<br>x10<sup>9</sup>/L | Monocyte<br>x10<sup>9</sup>/L | Eosinophil<br>x10<sup>9</sup>/L |
| --- | --- | --- | --- | --- | --- | --- | --- |
| *Erg<sup>fl/fl</sup>*<br>(n= 32) | 11.03 ± 0.48 | 1038 ± 192 | 9.07 ± 1.64 | 1.00 ± 0.90 | 7.52 ± 1.66 | 0.18 ± 0.15 | 0.21 ± 0.11 |
| *Rag1Cre<sup>T/+</sup>;Erg<sup>Δ/Δ</sup>*<br>(n=25) | 10.73 ± 1.85 | 1256 ± 204 | 4.82 ± 1.72 * | 1.09 ± 0.82 | 3.22 ± 1.12 * | 0.22 ± 0.17 | 0.24 ± 0.06 |

Blood was collected into EDTA and differential cell counts performed using an ADVIA 120 Hematology System. RBC, red blood cells; WBC, white blood cells. \* *P*<sub>adj</sub> < 10<sup>-11</sup> by Student's two-tailed unpaired t-test corrected for multiple testing by Benjamini-Hochberg procedure to control for false discovery rate.

**Supplementary Table S3.**
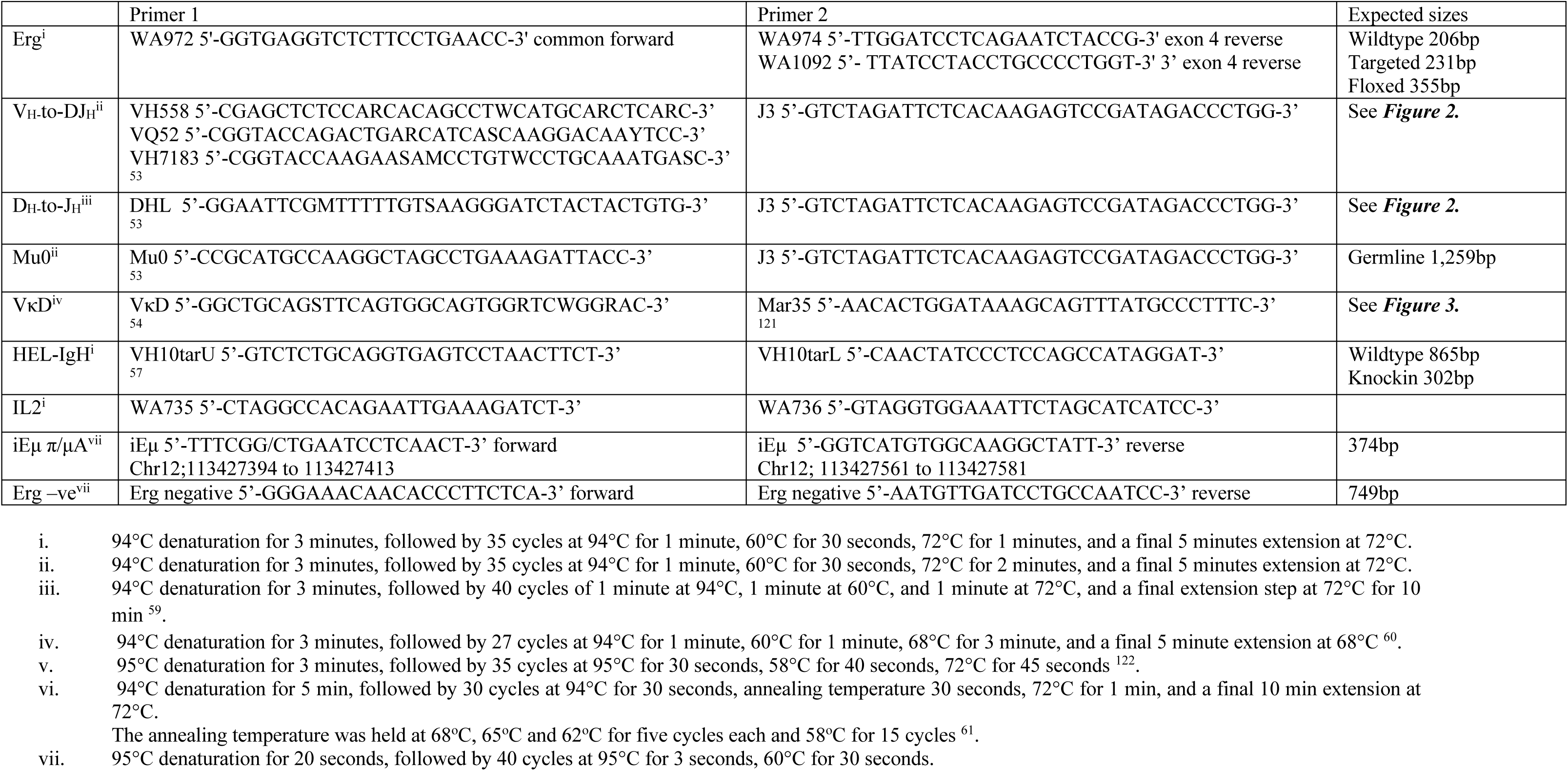
Primers and PCR reactions. See Figure 2, 3

## Notes

Conflict of Interest: **None**

